# Diverse long-range projections convey position information to the retrosplenial cortex

**DOI:** 10.1101/2022.09.18.508427

**Authors:** Michele Gianatti, Ann Christin Garvert, Koen Vervaeke

## Abstract

Neuronal signals encoding the animal’s position, originally discovered in the hippocampus, widely modulate neocortical processing. While it is assumed that these signals depend on hippocampal output, their origin has not been investigated directly. Here, we asked which brain region sends position information to the retrosplenial cortex (RSC), a key circuit for navigation and memory. Using two-photon axonal imaging in head-fixed mice performing a spatial task, we performed a comprehensive functional characterization of long-range inputs to agranular RSC. Surprisingly, most long-range pathways convey position information, but with key differences. We found that axons from the secondary motor cortex transmit the most position information. By contrast, axons from the posterior parietal-anterior cingulate- and orbitofrontal cortex and thalamus convey substantially less position information. Axons from the primary- and secondary visual cortex make a negligible contribution. These data show that RSC is a node in a widely distributed ensemble of networks that share position information in a projection-specific manner.

## Introduction

Hippocampal place cells are excitatory neurons that are active when an animal enters a specific location in its environment (O’Keefe and Dostrovsky, 1971). They likely form the neural basis of spatial memories by storing abstract relationships between sensory attributes experienced in a specific location (Eichenbaum, 2017; Robinson et al., 2020). It is increasingly evident that position-modulated spiking is also present in brain areas outside the hippocampus, where it is often multiplexed with sensory and motor information in a task-specific manner (Alexander and Nitz, 2015; Cho and Sharp, 2001; Esteves et al., 2020; Fiser et al., 2016; Harvey et al., 2012; Hok et al., 2005; Jankowski et al., 2015; Jankowski and O’Mara, 2015; Long and Zhang, 2021; Long et al., 2021; Mao et al., 2017; McNaughton et al., 1994; Mizumori et al., 1992; Nitz, 2006; Olson et al., 2020; Pakan et al., 2018; Quirk et al., 1992; Remondes and Wilson, 2013; Saleem et al., 2018). However, how the position code is generated outside the hippocampus, and what sources drive it, remains poorly understood.

Recently, position-tuned neurons reminiscent of hippocampal place cells were discovered in the retrosplenial cortex (RSC) (Campbell et al., 2021; Fischer et al., 2020; Mao et al., 2017), a major output structure of the hippocampus (Cembrowski et al., 2018; Kitanishi et al., 2021; Sugar et al., 2011; Vann et al., 2009; Wyss and Groen, 1992) and a brain area critical for storage and retrieval of spatial memories (Cowansage et al., 2014; Czajkowski et al., 2014; Milczarek et al., 2018; Vann and Aggleton, 2002). The origin of position signals to RSC is not known. Position information may be routed from the hippocampus to RSC via the subiculum (Ahmed and Mehta, 2009; Cembrowski et al., 2018; Kitanishi et al., 2021), supporting a model of the hippocampus as a central hub that instructs downstream circuits during memory retrieval (Teyler and DiScenna, 1986). Indeed, position tuning in RSC may depend, at least in part, on an intact hippocampus (Mao et al., 2018). However, there are also numerous pathways that support information flow in the opposite direction, from RSC to the hippocampus (Cenquizca and Swanson, 2007; Strange et al., 2014), and RSC may receive position-tuned information from other cortical or subcortical areas (Esteves et al., 2020; Mao et al., 2018).

An essential step toward determining the origin of the spatial signal in the cortex is to test whether neurons inherit their selectivity from tuned inputs originating from different upstream circuits. However, because measuring activity from the thin afferent axons in running mice and in a projection-specific manner is technically challenging (Broussard and Petreanu, 2021), a comprehensive overview of all input signals to a position-modulated circuit has not yet been achieved. Located by the brain’s surface, the agranular portion of RSC offers the unique opportunity to use two-photon Ca^2+^ imaging of long-range axons, enabling direct observation and characterization of the conveyed information (Petreanu et al., 2012). Here, we identified agranular RSC’s major presynaptic circuits using retrograde viral labelling (Tervo et al., 2016). Then, we trained head-fixed mice to perform a spatial task (Royer et al., 2012) and measured the activity of the projections from all major presynaptic circuits. Surprisingly, we found that most long-range inputs to agranular RSC convey position information, but in a projection-specific manner. These data show that RSC is a node in a widely distributed ensemble of networks that share position information.

## Results

### Functional imaging of long-range inputs to agranular RSC

We trained head-fixed mice to run laps on a circular treadmill to find the location of a water reward (26 mice, 66 sessions, 63 ± 15 laps per session, mean ± SD) (Figure 1A). Well-trained mice ran at a relatively constant speed but slowed down to lick for water at the same location, indicating that they learned the rewarded position (Figure 1B,C). This task resembles the one used in a recent study showing that up to 56 % of cells in layer 2/3 of agranular RSC cells have spatial tuning properties reminiscent of hippocampal place cells (Mao et al., 2017). The critical difference in our task was that mice ran in complete darkness to avoid visual responses. For this reason, we first needed to confirm that position tuned cells are present in agranular RSC under our task conditions. To measure neuronal activity, we performed two-photon Ca^2+^ imaging using mice expressing the Ca^2+^ indicator GCaMP6s in excitatory neurons (Thy1-GCaMP6s, 3 mice, 9 Fields of View (FOV), 1742 cells, see Methods) (Dana et al., 2014) and we deconvolved the fluorescence signals to infer spiking activity (Figure S1A, see Methods). Because RSC extends over several millimeters along the anterior-posterior axis, indicating heterogeneous connections and functions (Franco and Goard, 2021; Powell et al., 2020), we focused our analysis on L2/3 cells in the middle one-third of RSC where position-tuned cells were previously found (Mao et al., 2017). To analyze neuronal responses, we fitted linear-nonlinear-Poisson (LNP) models to the activity of each cell (see Methods) (Hardcastle et al., 2017). This enabled us to distinguish whether neurons respond to a specific position, or to sensory and motor variables such as tactile cues, running speed, acceleration, and lick rate. Neurons tuned to these sensory and motor variables may be confused with position-tuned cells if these variables are highly correlated with a specific location (see Methods). When mice performed this task in darkness, we found that a high fraction of cells in L2/3 of agranular RSC were position-tuned (33.1 ± 5.2 %, across 9 FOVs, 3 mice, (Figure S1B-D). Tuning properties of classified cells were stable throughout a session as this was an implicit criterion of the LNP model. Position tuned cells tiled the whole track, and using Bayesian decoding of neuronal activity (see Methods), we could reconstruct the mouse’s position with high precision (Figure S1E).

**Figure 1.**
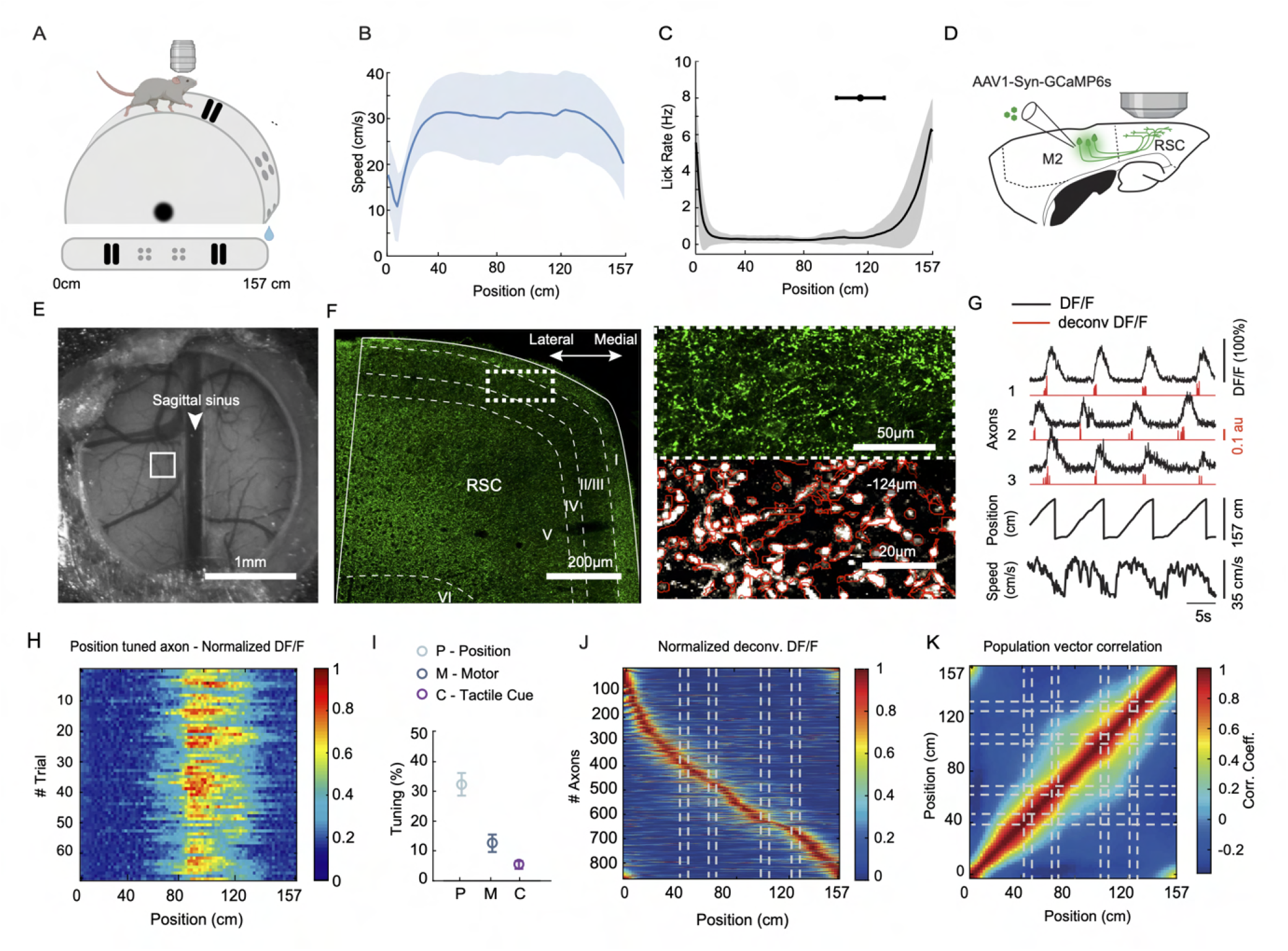
Functional imaging of M2 inputs to agranular RSC. (**A**) Behavior setup with a head-fixed mouse running on a Styrofoam wheel enriched with tactile cues (sandpaper strips, felt pads). Mice ran in complete darkness and received a water reward at the end of each lap. (**B**) Running speed profile of trained mice (mean ± SD across 66 sessions, 26 mice). (**C**) Lick rate of trained mice (mean ± SD). Horizontal Error bar shows the average location of the first lick before the reward (mean ± SD). (**D**) Diagram of the experiment. M2 neurons were infected with GCaMP6s expressing virus and their axons were imaged in L2/3 of agranular RSC. (**E**) Bright field image of the vasculature and implanted glass window above RSC. Square box shows typical field of view size and location for two-photon imaging. (**F**) (Left) Confocal fluorescence image of a coronal brain slice showing GCaMP6s-labelled M2 axons in RSC. Square box shows the typical location for two-photon imaging. (Right top) Expanded square box showing GCaMP6s-labelled M2 axons in deep layer 1 and layer 2/3. (Right bottom) Two-photon fluorescence image showing M2 axons and boutons (regions of interest in red). (**G**) Activity (DF/F) of three example M2 axons across four laps, aligned to position and running speed. Inferred spike rate (deconvolved DF/F) shown in red. (**H**) Activity (DF/F) of a position tuned M2 axon across a whole session. (**I**) Percentage of M2 axons tuned to different behavioral variables (mean ± SEM across 8 sessions, 4147 axons from 4 mice). “Motor” tuning included all cells tuned to running speed, acceleration, or lick rate. (**J**) Session-averaged activity of all position tuned M2 axons sorted by the location of peak activity. Dashed lines indicate tactile cue positions. (**K**) Correlation matrix (Pearson coefficient) of population activity vectors for all position-tuned axons (based on data in (J))

Next, we tested whether long-range projections convey position information to layer 2/3 of agranular RSC. We first focused on axons originating from the secondary motor cortex (M2) because this is a well-known dense projection to RSC (Yamawaki et al., 2016), and recent work found position tuned cells in M2 (Esteves et al., 2020). To image the activity of M2 axons in RSC, we expressed GCaMP6s in M2 neurons and implanted a chronic window over RSC (Figure 1D-E, 4 mice). Fluorescent axons and synaptic terminals were abundant in L2/3 of agranular RSC (Figure 1F). Because fluorescence signals of the same axon were highly correlated, we could identify boutons of the same axon, even when the axonal arborization could not be reconstructed morphologically (see Methods). This enabled us to measure activity from unique axons (4147 axons, 8 FOVs). Axonal imaging revealed highly position correlated neuronal activity that was reliable across laps (Figure 1G,H). Based on the LNP model, position tuning was the dominant response feature (32.1 ± 3.7 % of axons), with a minority of axons responding to motor variables (12.4 ± 2.8 %, running speed/acceleration and lick rate) and tactile cues (5.2 ± 1.1 %) (Figure 1I). When considering the entire population, position tuned M2 axons tiled the whole track with narrowly tuned firing fields (Figure 1J). Furthermore, consistent with an orthogonal code for position, a key feature of the hippocampal place code, population activity was more correlated between nearby positions than between more distant ones (see Methods, (Figure 1K)) (Mao et al., 2017). Altogether, these data show that a substantial fraction of M2 axons convey position signals to RSC while mice run on a linear track in the dark.

### Origin-specific differences in the fraction of position tuned axons

We tested whether axons from other presynaptic circuits also convey position information. First, we used viral retrograde labeling to identify the presynaptic circuits and injected rAAV2-retro-GFP in the superficial layers of agranular RSC (Figure S2A, 3 mice, see Methods). We limited injections to the middle one-third of agranular RSC, where we had measured position tuned activity (see Figure S1). Consistent with data from other species (Groen and Wyss, 1992), the areas with the densest somatic labeling included (from anterior to posterior): the orbitofrontal cortex (OFC, Figure S2B), M2 (Figure S2C), anterior cingulate cortex (ACC, Figure S2D), the anteroventral and anteromedial nucleus of the thalamus (AV and AM, Figure S2E), the posterior parietal cortex (PPC, Figure S2F), and the medial portion of the secondary visual cortex (V2M, Figure S2G). We also found somatic labeling in the claustrum (Figure S2C) and very sparse labeling in the entorhinal cortex (Haugland et al., 2019), but as we did not intend to characterize all RSC inputs exhaustively, we did not further study the projections from these two areas.

Next, we injected an AAV to deliver GCaMP6s under the synapsin promoter in each presynaptic area (in separate mice), and we verified whether neurons project to the superficial layers of agranular RSC (Figure 2A injection sites are shown in Figures S3-S7). Using two-photon microscopy in vivo, we observed GCaMP6-expressing axons originating from each presynaptic area (Figure 2A bottom row). When we measured axonal activity in mice performing the spatial task, we were surprised to find position-tuned axons among most long-range pathways (Figure 2B,C). Using the LNP model to determine whether axons were tuned to position, motor variables, or tactile cues (Figure S8), we found that position tuning was the dominant feature of most input pathways. There were, however, important differences (Figure 2D, E). Similar to M2 (32.1 ± 3.7 %), a high percentage of PPC axons was position-tuned (41.6 ± 7 %, 4 mice), followed by ACC (19.8 ± 2.1 %, 4 mice), OFC (14.3 ± 3.3 %, 5 mice) and thalamus (8.9 ± 1.8 %, 5 mice; note that we could not transduce neurons in specific thalamic nuclei, hence the general term “thalamus”). The outlier was V2M, with significantly fewer tuned axons than in most other areas (1.3 ± 0.4 %, 5 mice)(Figure 2E). For all areas, the percentage of position modulated axons was higher than expected by chance (0.16 ± 0.01 %, see Methods). These long-range inputs also differed in how population activity is correlated between nearby and distant positions along the track (Figure S9). As observed for M2 axons (Figure 1K) and RSC cells (Mao et al., 2017), the population activity of PPC, ACC and thalamic input was consistent with an orthogonal code for position, while input from OFC and V2M was too sparse to be considered orthogonal (Figure S9). We explored the possibility that the difference in the fraction of tuned cells was due to differences in the signal to noise ratio of the axonal fluorescence signals. However, the signal to noise ratio (Figure S10A) and the size of each axonal region of interest, which influences the signal to noise ratio (Figure S10B), did not correlate with the percentage of tuned cells. Altogether, we conclude that several presynaptic areas, in particular M2 and PPC, convey position signals to agranular RSC in this task.

**Figure 2.**
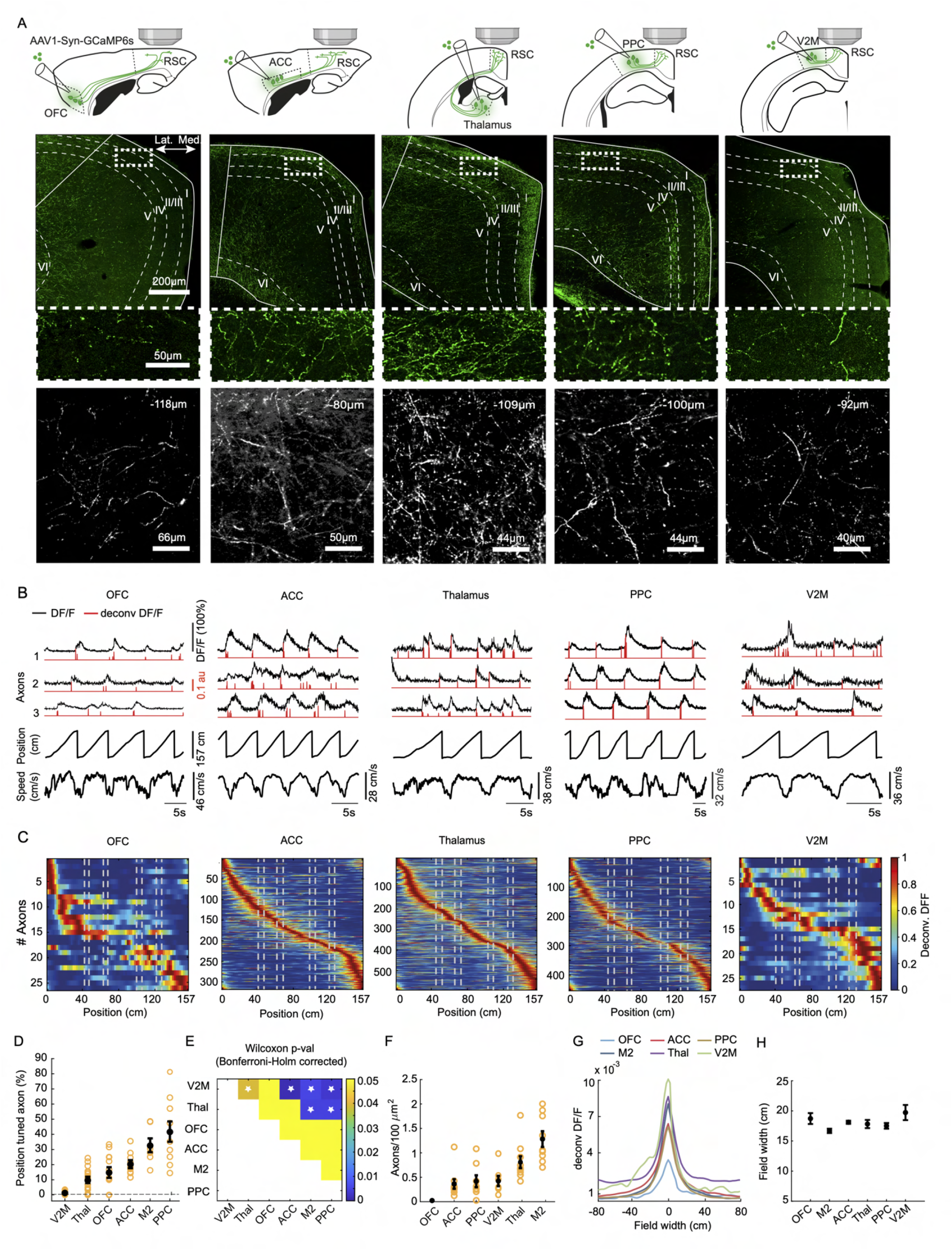
Origin-specific differences in the fraction of position tuned axons. (**A**)(Top row) Diagram of experiments. Neurons in indicated areas were infected with a GCaMP6s expressing virus and their axons were imaged in L1 and L2/3 of agranular RSC (OFC, orbitofrontal cortex; ACC, anterior cingulate cortex; PPC, posterior parietal cortex; V2M, medial secondary visual cortex). (Second row) Confocal fluorescence images of coronal brain slices showing GCaMP6s-labelled axons in RSC originating from the areas shown above. Square box shows the typical location for two-photon imaging. (Third row) Magnification of the square boxed area showing GCaMP6s-labelled axons in deep layer 1 and layer 2/3 of agranular RSC. (Bottom row) Two-photon fluorescence images showing typical fields of view with axons originating from the different areas. Imaging depth is indicated. (**B**) Activity (DF/F) of three example axons across 3-5 laps, aligned to position and running speed. Inferred spike rate (deconvolved DF/F) shown in red. (**C**) Session-averaged activity of all axons tuned to position and sorted by peak activity. Dashed lines indicate tactile cue positions. (**D**) Percentage of position tuned axons (mean ± SEM across sessions). The dashed baseline indicates chance level (V2M, 2918 axons from 5 mice, 13 sessions; Thalamus, 4536 axons from 5 mice, 18 sessions; OFC, 238 axons from 5 mice, 12 sessions; ACC, 2071 axons from 4 mice, 11 sessions; M2, 4147 axons from 4 mice, 8 sessions; PPC, 1516 axons from 4 mice, 9 sessions). (**E**) Statistical comparison of the percentage of position tuned axons between areas (Wilcoxon-Mann-Whitney test, Bonferroni-Holm multiple comparisons corrected). Stars indicate pairwise comparisons with p *<* 0.05. (**F**) Axonal density (mean ± SEM across sessions) from all areas based on two-photon imaging data of individual axons in agranular RSC.(**G**) Place field width and amplitude of position tuned axons (session-averaged deconvolved DF/F). (**H**) Place field width (mean ± SEM across sessions, full width at half maximum)

The low fraction of position tuned V2M axons was surprising because previous work reported large fractions of spatially modulated neurons in visual areas V1 and V2 measured using a similar task (Diamanti et al., 2021; Fischer et al., 2020; Fiser et al., 2016; Flossmann and Rochefort, 2021; Fournier et al., 2020; Haggerty and Ji, 2015; Pakan et al., 2018; Saleem et al., 2018). The key difference between these and our experiments is likely the lack of visual input in our task. We performed additional experiments and measured V1 axonal activity in the posterior part of RSC that receives more V1 input (Figure S11) (Groen and Wyss, 1992; Powell et al., 2020). However, the percentage of position tuned V1 axons was not above chance level (1.7 ± 1.6 %, 4 mice).

The influence of long-range input also depends on the density of their axons.Because we could measure fluorescence signals from individual axons, we could estimate the density of axons in the superficial layers of agranular RSC (Figure 2F). These data showed substantial differences in axonal density. OFC axons were very sparse, ACC, PPC and V2M intermediate, while M2 and thalamus axons had the highest density.

Neurons in the ventral hippocampus that project to the prefrontal cortex have place fields that are almost an order of magnitude larger than those of the dorsal hippocampal neurons that project indirectly to the retrosplenial cortex (Jay and Witter, 1991; Kjelstrup et al., 2008; Strange et al., 2014). Inspired by this observation, we hypothesized that axons in RSC, originating from ACC and OFC have larger place fields than axons from more posterior cortical areas. However, this was not the case (Figure 2G). The mean tuning field width was about 18 cm for all long-range inputs, closer to the place field width of neurons in the dorsal hippocampus (Kjelstrup et al., 2008) (Figure 2H). The highly similar place field width was not a consequence of how we classified tuned cells because our LNP model, unlike other classification methods, does not use tuning field width as a criterium.

Altogether, these data show significant origin-specific differences in the fraction of position tuned axons and axonal density in the superficial layers of agranular RSC. In contrast, the width of position tuning is strikingly similar for all long-range inputs.

### Origin-specific differences in spatial information conveyed to RSC

Next, we measured how much position information the various long-range projections convey to agranular RSC. Using Bayesian decoding of axonal activity, we quantified the difference between estimated and actual mouse position (see Methods). First, we used all axons in a field of view (FOV), including classified axons (position, tactile cues, and motor variables) and unclassified axons (Figure 3A,B). M2 axons had the lowest decoding error (10.8 ± 1.0 cm), followed by axons from the thalamus (20.9 ± 1.1 cm), PPC (22.5 ± 2.9 cm) and ACC (25.3 ± 1.5 cm) (Figure 3B). The decoding error was the highest for V2M (28.5 ± 1.1 cm) and OFC axons (36 ± 0.6 cm) (Figure 3B). A statistical comparison between projections showed that the decoding error using M2 axonal signals was significantly lower than most other projections (Figure 3C). Because the decoding error depends on the axons’ density, we subsampled the number of axons from each presynaptic area and quantified the decoding error as a function of axonal population size (Figure 3D). When we considered a similar number of axons from each area (270 axons), we found that decoding from M2 and PPC axons was more accurate than the other projections (Figure 3D, dashed box). Next, we decoded position using only position classified axons. Overall, most inputs’ decoding error increased slightly (Figure 3E). This suggests that excluded axons, such as those responding to sensory cues and motor variables, also contribute position information. Nevertheless, M2 (12.7 ± 1.3 cm) and PPC axons (21.7 ± 2.4 cm) performed the best, with M2 axons having a significantly lower decoding error than most other areas (Figure 3F). Interestingly, when we subsampled the axonal population from each area and considered a common number of axons (40 axons), the decoding error was similar for M2, PPC, AAC and thalamic input (Figure 3G, dashed box). This shows that M2 conveys more position information because position tuned M2 axons in agranular RSC are more numerous, not because the position information per M2 axon is higher than in other areas.

**Figure 3.**
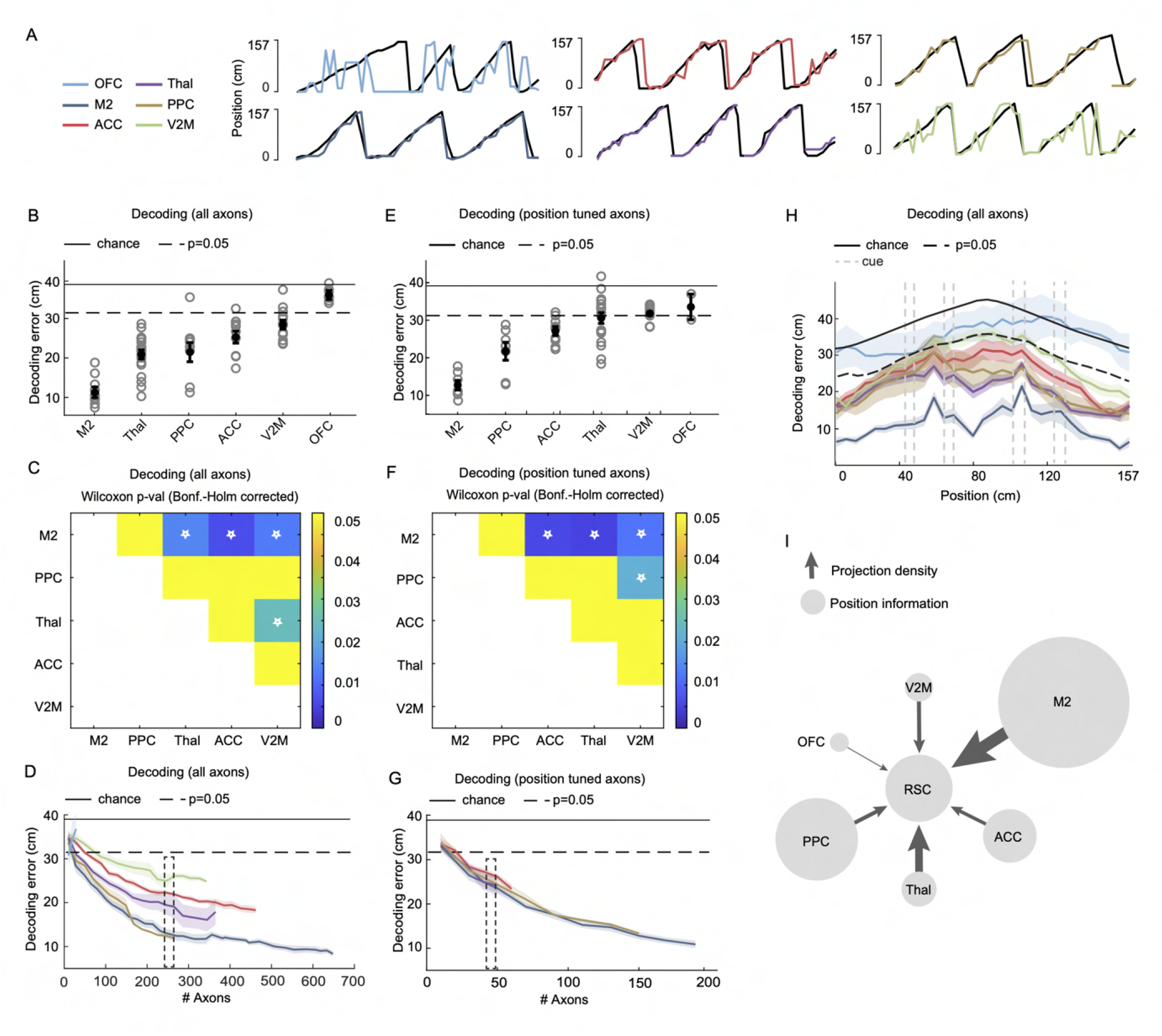
Origin-specific differences in spatial information conveyed to RSC. (**A**) Examples of actual (black) and estimated (color) mouse position obtained by Bayesian decoding using the activity of all recorded axons in a field of view (FOV). (**B**) Decoding error (mean ± SEM across sessions) using all recorded axons in a FOV (regardless whether classified or not). (**C**) Statistical comparison of decoding errors between areas (Wilcoxon-Mann-Whitney test, Bonferroni-Holm multiple comparisons corrected). Based on the data in(B). Asterisks indicate comparisons with p *<* 0.05. (**D**) Decoding error (mean ± SEM across sessions) as a function of number of axons used for decoding. Dashed box is centered on 270 axons. (**E**) Decoding error (mean ± SEM across sessions) using only position tuned axons in a FOV during a session. (**F**) Statistical comparison of decoding errors between areas (Wilcoxon-Mann-Whitney test, Bonferroni-Holm multiple comparison corrected). Based on the data in (E). Asterisks indicate pairwise comparisons with p *<* 0.05. (**G**) Decoding error as a function of the number of axons used for decoding (mean ± SEM across sessions). Only position tuned axons in a FOV were used. Dashed box is centered on 40 axons. (**H**) Decoding error (mean ± SEM across sessions) as a function of position using all available axons in a FOV (regardless whether classified or not). (**I**) Summary of long-range projections conveying position information to agranular RSC. Arrow thickness represents the projection density (see Figure 2F), and bubble size represents spatial information (see Figure 3E). The diameter of a bubble indicates the difference between an area’s decoding error and the chance-level decoding error.

We verified that these differences were not due to differences in running speed between mice. We found that mice with GCaMP6 labeled axons from different brain areas showed a similar running speed profile but different speed amplitude (Figure S12A, B). However, the decoding error and running speed amplitude did not correlate when considering all sessions of all mice (Figure S12C, see Methods). We also tested how the decoding error depends on the number of trials and whether this varied between mice. The decoding error only marginally relied on the number of trials (Figure S12D), and the number of trials per session was similar for all mice (Figure S12E).

Finally, we quantified the decoding error along the track using all classified and unclassified axons (Figure 3H). The decoding error is the lowest around the reward location and substantially higher in the middle of the track. Again, the decoding error using M2 axons was lower along the whole track than any other long-range input. We summarized these data in a bubble diagram using only position tuned axons (Figure 3I). Altogether, of all long-range pathways tested, M2 and, to a lesser degree, PPC stand out as the dominant source of position information, while V2M and OFC contributed negligibly.

## Discussion

To investigate how position tuning is generated in RSC, we performed a comprehensive functional analysis of its long-range inputs. We determined the areas sending long-range projections to agranular RSC and found that most of them transmit signals that enable decoding the mouse’s position with high precision. There were, however, significant differences between projections. M2 provided significantly more position information than any other area, followed by PPC, ACC, thalamus and OFC, while the visual areas V2M and V1 provided little or none.

Recent studies show that brain areas outside the hippocampus show position modulated activity. Such extra-hippocampal spatial signals may be important for contextualizing sensory and motor signals experienced in different locations (Flossmann and Rochefort, 2021; Teyler and DiScenna, 1986). However, in some cases this activity does not unequivocally encode position but may represent location-specific sensory cues, motor signals or rewards (Grieves and Jeffery, 2016; Knierim, 2006; Peyrache and Duszkiewicz, 2021). For this reason, we performed experiments in total darkness, and we used the LNP model to test whether position-tuned RSC neurons and their afferent inputs respond to tactile cues or motor variables such as speed, acceleration, or lick rate (Hardcastle et al., 2017). After applying these criteria, we found that several long-range projections have a remarkably high fraction of position-tuned axons, particularly M2 and PPC.

Long-range projections from V1 and V2M transmitted little or no position information to RSC. Given the abundance of recent work showing position modulated activity in V1 and V2, this result was surprising (Campbell et al., 2021; Diamanti et al., 2021; Fischer et al., 2020; Fiser et al., 2016; Flossmann and Rochefort, 2021; Haggerty and Ji, 2015; Pakan et al., 2018; Saleem et al., 2018). However, in contrast to previous work, our experiments were performed in darkness to avoid visual input. Therefore, our data suggest that spatial modulation in V1 cannot be maintained by self-motion or other sensory cues in the absence of visual input. Alternatively, spatial signals in V1 may be primarily visual signals modulated by position.

Head-fixed tasks on a linear track offer important advantages for studying circuit mechanisms. However, our spatial task only tests an egocentric search strategy called path integration (Campbell et al., 2021; Mao et al., 2020). The animal deduces position by estimating distance from a reference point (here, likely the reward location) based on self-motion information (McNaughton et al., 2006). However, hippocampal place cells also provide an allocentric (world-centered) code for position in freely moving animals exploring a two-dimensional arena (Wang et al., 2020). Such an allocentric code has not been unequivocally established in circuits outside the hippocampus (Grieves and Jeffery, 2016; Knierim, 2006; Peyrache and Duszkiewicz, 2021). Therefore, it remains to be determined whether our observations also apply to an allocentric spatial code.

The distributed nature of position signals raises the question of where position tuning is generated. Previous work suggested that CA1 hippocampal neurons inherit their spatial firing patterns from area CA3 and entorhinal cortex and route this to output structures such as RSC and the medial entorhinal cortex (Ahmed and Mehta, 2009; Cembrowski et al., 2018; Kitanishi et al., 2021; Skelin et al., 2019). RSC could thus act as a hub for distributing spatial information to neocortical circuits (Nitzan et al., 2020; Teyler and DiScenna, 1986). Here, we did not record from hippocampal inputs in agranular RSC because retrograde labeling did not provide evidence for a direct connection. A hippocampal position signal may reach agranular RSC indirectly via the subiculum that terminates in the granular portion of RSC, which in turn projects to agranular RSC. Instead, we find that RSC receives direct spatial input from several neocortical circuits and thalamic nuclei. This could be explained by the reciprocal connections between neocortical circuits that enable a bidirectional exchange of position signals. Indeed, such reciprocal connections exist between RSC and M2, both monosynaptically, and disynaptically via PPC and AM thalamus (Yamawaki et al., 2016). However, we cannot exclude the possibility that areas outside the hippocampus can also generate position tuning. In support of this, a recent theoretical study showed that circuits could generate neuronal sequences, reminiscent of position tuning, with only a simple learning rule and without the need for highly specific connections (Rajan et al., 2016). This challenges the idea of the hippocampus as the main driver of position signals in the cortex. Therefore, future experiments will need to address whether neocortical circuits can also generate position-tuned responses, independently from the hippocampus, and how these circuits interact during navigation, memory formation and memory retrieval.

## ACKNOWLEDGEMENTS

This work was funded by the European Research Council (ERC Starting Grant #639272), the Research Council of Norway (#274306), an EEA grant (RO-NO-2019-0504), and a PhD position from the University of Oslo. We thank Kristin Larsen Sand, and Eivind Hennestad for technical assistance, Menno Witter for help with neuroanatomy and delineations, and Aree Witoelar and Benjamin Dunn for mentoring ACG. We thank Bruno Pichler (Independent NeuroScience Services; INSS) for developing the custom two-photon microscope. Viral constructs were gifts from Douglas Kim & GENIE project. Some schematics were created with Biorender.com. We thank Matthijs Dorst, Torkel Hafting, Hua Hu, Jørgen Sugar, Jonathan Whitlock, Menno Witter and members of the Vervaeke lab for providing comments on a draft of the manuscript.

## AUTHOR CONTRIBUTIONS

MG, ACG and KV designed the study. MG performed all experiments and non-model-based analysis. ACG performed all model-based analyses. MG, ACG and KV wrote the paper. All authors discussed the results.

## Methods

### Resource availability

#### Lead contact

Further information and resource requests should be directed to and will be fulfilled by the lead contact Koen Vervaeke.

#### Materials availability

This study did not generate new unique reagents.

#### Data and code availability

- Ca^2+^ imaging and tracing data reported in this paper will be shared by the lead contact upon request.
- All original code will be deposited on GitHub and is publicly available as of the date of publication. Any additional information required to reanalyze the data reported in this paper is available from the lead contact upon request.

### Experimental model and subject details

For axonal calcium imaging, we used 31 adult female mice (18-24g, C57Bl/6J; Janvier labs, 2-4 months old at surgery). For somatic calcium imaging, we used 3 Thy1-GCaMP6s male adult mice (20-24g, GP4.3 line, Jackson Laboratory #024275, 2-4 months old at surgery). For retrograde tracing experiments, we used 3 female mice (18-24g, C57Bl/6J; Janvier labs, 2-4 months old at surgery). We kept mice on a reversed 12-hour light/ 12-hour dark cycle for behavioural experiments, and we performed experiments during the dark phase. Mice received water drop rewards throughout the task, and we kept them on a water restriction regime as described in (Guo et al., 2014). All procedures were approved by the Norwegian Food Safety Authority(FOTS #19129). Experiments were performed in accordance with the Norwegian Animal Welfare Act.

### Methods details

#### Experimental setup

The running wheel was a polystyrene cylinder with a circumference of 157 cm circumference and a width of 10 cm (Bakedeco, Brooklyn, New York). There were two types of tactile cues, black sandpaper strips (10 cm wide, 2,5 cm long) and white soft felt pads (also called furniture pads, 2 cm diameter, 3 mm high). These were glued at two positions on the wheel; 42-47 cm and 122-128 cm (sand-paper strips) and 63-68 cm and 100-106 cm (felt pads) (the layout is shown in Figure 1B). To fix the mouse’s head, we used a head post and clamp system (Guo et al., 2014). We controlled water reward delivery with a solenoid valve and a custom-made lick port. The lick port was part of a custom-built electrical circuit to detect when the tongue contacted the lick port (Guo et al., 2014). We calculated running speed and absolute position on the wheel using a rotary encoder. We conducted all experiments in total darkness in a sealed box consisting of black construction hardboard (Thorlabs, TB4) and black coated fabric (Thorlabs, BK5).

#### Behavior control and acquisition software

We automated the behavioral task through a custom-written LabVIEW program (LabVIEW 2013, National Instruments). The lick-port signals, rotary encoder signals and two-photon frame-clock were acquired using a DAQ (X-Series, PCIe 6351, National Instruments). We used the two-photon frame clock signal to synchronize the two-photon images with the recorded signals via the DAQ.

#### Surgery

Surgery was carried out under isoflurane anesthesia (3 % induction, 1-1.5 % maintenance) while maintaining body temperature at 37°C with a heating pad (Harvard Apparatus). We delivered a subcutaneous injection of 0.1 mL Marcaine (bupivacaine 0.25% m/V in sterile water) at the scalp incision site and administered post-operative analgesia (Temgesic, 0.1 mg/kg) subcutaneously. Several weeks before experiments began, we implanted the head bar and cranial window. The center of the window was −2.2 mm AP from bregma. The window consisted of a circular outer window (3.5 mm diameter, #1.5 coverslip glass) affixed to a circular inner window (2.5 mm diameter, #2 coverslip glass) with optical adhesive (Norland Optical Adhesive; ThorLabs NOA61). We held the window in place by applying gentle pressure so that the outer window would fit into the thinned skull area and flush with the skull’s surface. We used heated agar (1 %; Sigma #A6877) to seal any open spaces between the skull and edges of the glass window, and we affixed the window to the skull with cyanoacrylate glue. A detailed protocol of the surgery can be found in (Holtmaat et al., 2009).

#### Water restriction

Starting at least 1 week after surgery, we placed mice on water restriction (Guo et al., 2014). Mice received 1 − 1.5 ml of water per day while we monitored their body weight to ensure they maintained more than 80 % of the initial body weight.

#### Animal training / habituation

During the first week of water restriction, we handled mice daily in semi-darkness to make them comfortable with the experimenter. To habituate mice to head fixation and run on the wheel, we placed them gently on the wheel and let them run freely for 3 min. During training, mice first learned to run on a blank polystyrene wheel, and we gradually introduced the tactile cues. We trained mice for 1-3 weeks, or until they performed *>*100 laps in 20 min. Mice always received 2 *μ*l water drops at a fixed position on the wheel. We moved the setup under the microscope for data acquisition when the mice reached the criteria.

#### Two-photon microscopy

We used a custom-built two-photon microscope (INSS), designed to provide enough space under the objective to accommodate the large running wheel. We acquired images at 31 Hz (512 × 512 pixels) using SciScan (opensource, LabVIEW, National Instruments). The excitation wavelength was 950 nm using a MaiTai DeepSee ePH DS laser (SpectraPhysics). The average power measured under the objective (N16XLWD-PF, Nikon) was 50-100 mW. Photons were detected using GaAsP photomultiplier tubes (PMT2101/M, Thorlabs). The primary dichroic mirror was a 700 nm LP (Chroma), and the photon detection path consisted of a 680 nm SP filter (Chroma), a 565 nm LP dichroic mirror (Chroma), and a 510/80 BP filter (Chroma). The FOV size varied between 182-400*μ*m for axonal imaging and 454-666 *μ*m for somatic imaging. We recorded axons in deep L1 and superficial L2/3 of agranular RSC (70-130 *μ*m depth from the pial surface). We performed somatic recordings at 100-120*μ*m below the pial surface, corresponding to L2/3 of agranular RSC. Because imaging of subcellular structures like axons in running mice causes substantial brain motion artefacts, we stretched the point spread function (PSF) of the excitation volume by underfilling the objective back aperture (Broussard and Petreanu, 2021; Petreanu et al., 2012; Zipfel et al., 2003). To achieve an axial PSF of ∼5 *μ*m, we bypassed the telescope, leaving a 5 × beam magnification by the scan and tube lens assembly (see Figure S13E) for PSF measurements).

#### Viruses and retrograde tracing procedure

We injected a single 100 nL of AAV2-retro-GFP (Addgene #37825-AAVrg) in the left agranular RSC (AP: −2.18 mm; ML: 0.5 mm; DV: −0.2 mm) of C57Bl/6J mice. Mice expressed the virus for 4-5 weeks before perfusion and brain sectioning for histology. In two additional C57Bl/6J mice, we performed two 100 nL injections, offset by 1 mm AP, to express two fluorescent proteins with different colors in presynaptic areas (AAV2-retro-GFP and pAAV2-retro-tdTomato, Addgene # 59462). The aim was to test whether the labeled presynaptic circuits would differ substantially if the injection location was slightly jittered, which was not the case (AP: −1.8; ML 0.4; DV: −0.15 and AP −2.7; ML: 0.45; DV: 0.25 respectively).

For imaging experiments, we injected AAV1-Syn-GCaMP6s-WPRESV40 (Addgene #100843-AAV1) in the left hemisphere. Target coordinates were based on the retrograde labeling experiments: OFC (5 mice; AP: +3.48 mm; ML: 1.25 mm; DV −1.95 mm; volume 15-20 nl), ACC (4 mice; AP: +1.1 mm; ML: 0.30 mm; DV: −1.5 mm; volume 25 nl), M2 (4 mice; 2 injections, AP: −0.1 mm; ML: 0.5 mm; DV: −0.5 mm and AP: −0.1 mm; ML: 0.6 mm; DV: −0.2 mm; volume 20-25 nl each); thalamus (5 mice; AP: −1.22 mm; ML: 1.10 mm; DV: −2.60 mm; volume 30-40 nl); PPC (4 mice; AP: −1.95 mm; ML: 1.4 mm; DV: −0.5 mm and −0.2 mm; volume 15-20 nl each); V2M (4 mice; AP: −2.7 mm; ML: 1.5 mm; DV: −0.3 mm; volume 15 nl), V1 (4 mice; 2 injections, AP: −3.57 mm; ML 2.49 mm; DV: −0.3 mm and −0.5 mm; 20-30 nl).

#### Tissue collection and digitalization

We deeply anesthetized mice with pentobarbital sodium (90 mg/kg) and, once the reflexes were absent, transcardially perfused with 4 % paraformaldehyde (w/v) in phosphate-buffered saline (PBS). We kept the brains for at least 24 hours in 4 % paraformaldehyde, then transferred them to a cryoprotective solution (2% DMSO in 0.125M phosphate buffer) the next day and stored them overnight at 4 °C. We then embedded the brains, froze them at −40 °C and cut them into 40 *μ*m coronal sections with a freezing microtome. We collected three sets of sections; the first section was mounted while each 2nd and 3rd section was stored in cryoprotective solution at −20 °C, either for later GFP immunohistochemistry or for backup. We mounted the sections directly onto Superfrost Plus microscope slides (Gerhard Menzel GmbH, Braunschweig, Germany), dried them overnight on a heating pad, and then cleared them in xylene, and coverslipped them with Eukitt (Sigma-Aldrich). We then performed fluorescence imaging of the sections with a Zeiss Axio Scan.Z1 scanner. Next, we took off the coverslip and stained the same sections with cresyl violet (Sigma-Aldrich) to facilitate the delineation of injection sites. Briefly, we dehydrated the sections in increasing percentages of ethanol (50 %, 70 %, 80 %, 90 %, 3 × 100 %, 10 dips each). We briefly rinsed the sections in running water before staining them with cresyl violet (0.1 %) on a shaker for 3 min. To achieve optimal staining, we subsequently rinsed the sections in running water and differentiated them in an ethanol-acetic acid solution (0.5 % acetic acid in 70 % ethanol). We then dehydrated the sections again in increasing percentages of ethanol (as described above), cleared them in xylene, and coverslipped them with Eukitt (Sigma-Aldrich). We then imaged the Nissl-stained sections using a Zeiss Axio Scan.Z1 scanner.

#### Analysis of injection sites

To help delineate the injection sites, we superimposed the fluorescence and Nissl images and adjusted their contrast and brightness in Adobe Photoshop. We delineated all areas following (Paxinos and Franklin, 2019), except for PPC that was delineated following (Hovde et al., 2018).

#### GFP Immunohistochemistry

To enhance the fluorescence signal from GCaMP6s-positive axonal terminals in agranular RSC, we counterstained sections with antibodies against GFP. We first rinsed brain sections 3 × 5min in PBS on a shaker, incubated them in blocking buffer (PBS plus 0.3 % Triton, 2 × 10 min), and then incubated them in primary antibody solution (rabbit anti-GFP, 1:1000, ThermoFisher Scientific, A-11122, in PBS and 0.3% Triton) overnight at 4 °C. We further washed sections in PBS containing 0.3 % Triton and 3 % goat serum (Sigma Aldrich) for 2 × 5 min at room temperature (RT). Subsequently, we incubated them in secondary antibody solution (AlexaFluor 488-tagged goat anti-rabbit Ab, 1:1000, ThermoFisher Scientific, A-11008) for 1h at RT. We washed sections 2×10 min in PBS, mounted them onto Superfrost Plus microscope slides (Gerhard Menzel GmbH, Braunschweig, Germany), dried them overnight on a heating pad and then cleared them in xylene, and cover-slipped them with Eukitt (Sigma-Aldrich). Then, we scanned sections with a Zeiss confocal microscope. Finally, we Nissl-stained the sections using the same procedure described earlier and digitized them using a Zeiss Axio Scan.Z1 scanner. We then delineated the layers in agranular RSC according to (Sugar et al., 2011) in Adobe Illustrator CC 2021 by overlaying the fluorescent sections with their corresponding Nissl-stained sections.

### Quantification and statistical analysis

#### Image registration and segmentation

To correct brain movement, we registered images using a combination of custom-written Matlab scripts (NANSEN) and NoRMCorre (Pnevmatikakis et al., 2016). We segmented registered images using custom-written MATLAB scripts (NANSEN). We detected soma regions of interest (ROI) and axonal ROIs using a custom auto-segmentation method (NANSEN) followed by manual curation after visual inspection of each ROI. When ROIs were overlapping, the overlapping parts were excluded. For neuropil correction of the axonal signals (see section below: Signal and event extraction), a doughnut-shaped ROI of the surrounding neuropil was automatically created for each ROI, by dilating the ROI so that the doughnut area was four times larger than the soma/axon ROI area. If this doughnut overlapped with another soma/axon ROI, then that ROI was excluded from the doughnut.

#### Clustering method to detect individual axons

To avoid multiple ROIs of the same axon, we assigned the activity of ROIs to unique axons. We found ROIs belonging to the same axon by applying a hierarchical clustering algorithm (Murtagh and Contreras, 2012) using the MATLAB Statistics and Machine Learning Toolbox functions pdist(), linkage() and cluster(). As a starting point, we defined the similarity of the time series a and b of two ROIs with n ∈ N measurements as the Pearson’s correlation coefficient:

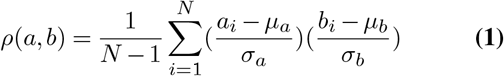

With *μ*_*a*_ and *σ*_*a*_ the mean and standard deviation of a, and *μ*_*b*_ and *σ*_*b*_ being the mean and standard deviation of b. We calculated the dissimilarity of a and b as:

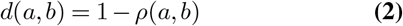

for every pair of ROIs with the MATLAB function pdist(). We then used the MATLAB function linkage() to sequentially find the two clusters with the greatest similarity (smallest dissimilarity) and merge them into one cluster. Initially, we treated each ROI as a separate cluster. We defined the dissimilarity D(A,B) between two clusters A and B containing multiple ROIs as the average of all pairwise dissimilarities between all ROI signals of the respective two clusters:

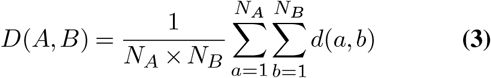

With *N*_*A*_, *N*_*B*_, the number of ROIs of cluster A and B. We created a hierarchical cluster tree by continuous merging until only one cluster containing all ROIs remained. At the end we pruned the tree to divide the ROIs into clusters using the MATLAB function cluster() for which we set a similarity threshold of 0.7 (dissimilarity of 0.3). The resulting clusters, consisting of multiple ROIs, were defined as unique axons.

#### Signal and event extraction

First, we averaged the pixel values of each ROI per imaging frame. We then defined ROI fluorescence changes as a fractional change DF/F(t) = (F(t)-F0)/F0, with F0 being the baseline defined as the 20th percentile of the ROI fluorescence signal F(t). We calculated this for soma/axon and neuropil ROIs. We then subtracted the neuropil DF/F from the soma/axon DF/F, and we added a correction factor to ensure that the soma/axon DF/F remained positive. We then deconvolved the resulting DF/F signal using the CaImAn package (Friedrich et al., 2017; Giovannucci et al., 2019) to obtain event rates that approximate the cell’s activity level.

#### Calculation of position tuning curves

We binned the deconvolved DF/F for each axon in 2 cm spatial bins. We then convolved this vector with a Gaussian window (4 cm standard deviation). Finally, we created the position tuning curve by averaging these activity vectors across all trials.

#### Correlation matrix of population activity vectors

To test the orthogonality of the axonal population activity vectors, we quantified the response similarity between axons tuned to distinct locations on the track. We calculated the position tuning curves for all position tuned axons and the pairwise Pearson correlation coefficients between the vectors of axonal activities for each position.

#### Classification of cue tuned cells

We calculated position tuning curves for each axon and detected all peaks higher than 30 % of the highest peak using the MATLAB Signal Processing Toolbox function findpeaks(). Then we required that the activity within the detected field around the peak is at least three times higher than the activity outside these fields. Finally, a reliability criterion required that the activity peak was present in these fields in at least 20 % of trials. Then we excluded peaks around the reward location (*<* 40 cm, *>* 140 cm on the track). Axons were classified as cue tuned if the distance between two fields was similar to the distance (± 6 cm) between the center position of two similar cues. Axons classified as cue tuned were not further tested for other covariates.

#### Classification of whether axons are tuned to position, running speed, lick rate or acceleration

We fitted 4 linear-nonlinear-Poisson (LNP) models (Hardcastle et al., 2017) for each axon using one of the following covariates: position (P), running speed (S), lick rate (L) and acceleration (A), and we identified which model fitted the data best.

We prepared the fitting process by first converting each of the covariates C = [P, S, L, A] into *N*_*c*_ bins. We then created a state vector *X*^*c*^ for each covariate in which each column corresponded to a time *t* ∈ *T* and each row corresponded to a bin of parameters *n*_*c*_ ∈ *N*_*c*_. The state matrix was set to *X*^*c*^(*n*_*c*_, *t*) = 1 for the parameter bin *n*_*c*_ that the animal experienced at time *t* and *X*^*c*^(*n*_*c*_, *t*) = 0 otherwise.

For a given model *M* ^*c*^ we assumed that the probability of recording the activity *k*_*t*_ in time bin *t* of length *dt* (*dt* = 32,3 ms) follows a Poisson distribution:

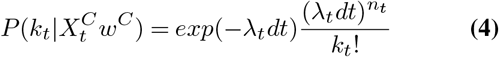

The variable parameters *w*^*C*^ convert the state vector *X*^*C*^ for covariate C into an expected firing rate *λ*_*t*_:

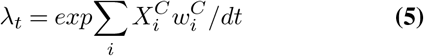

where i indexes over parameter elements for the covariate *C*. For every observed axonal activity train *k*_*t*_, we learned the parameters *w*^*C*^ for each of the models *M* ^*c*^ by optimizing the cost function (using MATLAB’s fminunc function):

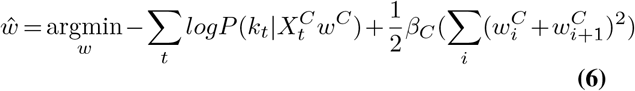

Whereby we impose smooth parameters with *β*_*c*_, the constant smoothing hyperparameter and trained the model in a 5-fold cross-validation procedure.

The tuning to each of the covariates was finally defined as the difference between the log likelihood of the model containing the covariate information and the log likelihood of the model with a fixed, mean firing rate:

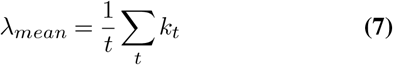

With T, the number of time samples.

Hyperparameters such as the bin numbers *N*_*c*_ of the covariates and the smoothing parameters were optimized over the entire population of axons for several sessions for each area (example shown in Figure S14A). The hyperparameters leading to the greatest log likelihood difference were chosen and did not differ between areas and adding more sessions did not change them. This optimization resulted in *N*_*p*_= 40 (3.925 cm bins width), *N*_*S*_ = 30 speed bins (2 cm/s bin width), *N*_*L*_= 20 bins lick rate bins (0.5 Hz bin width) and *N*_*A*_= 40 acceleration bins (10 *cm/s*^2^ bin width). The optimized smoothing hyperparameters were *β*_*P*_ = 0.01,*β*_*S*_ = 0.01,*β*_*L*_= 0 and *β*_*A*_= 0.

#### Characterization of tuning properties

We selected all models that performed significantly better than a fixed mean firing rate model to determine which of the variables (P, S, L or A) the axons encode for. We determined significance through a one-sided signed rank test where we tested the model’s log-likelihood containing the covariate information against the model’s log likelihood using a fixed mean firing rate of the 5 cross validation folds with significance value of p = 0.1. We classified axons for which no model performed better than the fixed mean firing rate model as not tuned (examples are shown in Figure S14B-H).

If several models passed this criterium, we tested if one model performed significantly better than the others. If the performance of one model was not significantly better than another, axons were defined as tuned to several covariates. To estimate the percentage of classified axons expected by chance, we randomly permuted the covariates in relation to the neural activity and applied the same classification procedure for each axon of each session. Chance levels were 0.16 ± 0.01 % position tuned cells, 0.03 ± 0.13 % running speed cells, 0 ± 0 % acceleration tuned cells, 0.09 ± 0.01 % lick rate tuned cells and 0.004 ± 0.00 % tactile cue tuned cells (mean ± SEM across sessions of all areas).

#### Bayesian decoding of position

The probability of an animals’ position is given by:

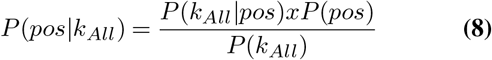

Assuming statistical independence between axonal signals *P*(*k*_*All*|*pos*_) can be written as:

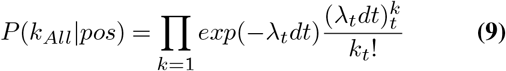

where k indexes over all axonal signals and the firing rates *λ*_*t*_.

We estimated the parameters *λ*_*t*_ as described and optimized the hyperparameters to minimize the decoding error. We binned the session data non-overlapping time bins (3 time bins 0.1 s), separated the positions on the wheel into 30 evenly sized bins (5.233 cm) and set the smoothing parameter to *β*_*P*_ = 0.01.

We used a flat prior P(pos), with which we assumed the animals do not have a certain expectance about the occupancy of the positions on the wheel, and the probability for each position at time point t is given by:

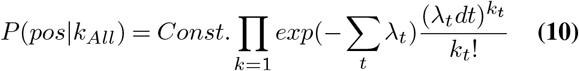

With Const. being a normalization factor so that *P* (*pos|k*_*All*_) sums to 1. We defined the position with the highest (max) probability as the decoded position for time point t.

We quantified the decoding error as the mean distance between predicted and actual position on the wheel. We trained the model in a 5-fold cross-validation procedure and defined the fold that led to the median decoding error as the resulting model.

We calculated the decoding error with varying trial numbers and concluded that already 30 trials are enough to reach a saturated decoding error.

To calculate the chance level for decoding we randomly shuffled the predicted position against the actual position 5000 times. We repeated this procedure for all sessions and for the decoding using different numbers of axons. We calculated the decoding error as described for these shuffled pairs and the final chance and p-value level of 0.05 was determined by the mean and the 5th percentile of these decoding errors.

#### How does running speed affect the decoding error?

To test whether the running speed of the mouse impacts the decoding results, we simulated data based on two different speed profiles of real sessions. We choose one session in which the mouse was particularly slow (average speed: 16.9 cm/s) and another in which the mouse was running fast (average speed: 46.7 cm/s). For both sessions we generated respectively 31 spike trains *k*_*t*_ assuming a Poisson process. Hereby the firing rates *λ*_*t*_ were position dependent, imitating the firing of position cells through a gaussian. We chose the means of the gaussian distributions of firing rates so that all positions were evenly covered (means were about 5 cm distanced) and generated random values between 1 and 30 cm as the standard deviations that determined the width of the position fields, which is similar to what we observed in real data.

Next, we approximated calcium signals by convolving the spike trains with a fast rising and exponentially decaying function (Lütcke et al., 2013):

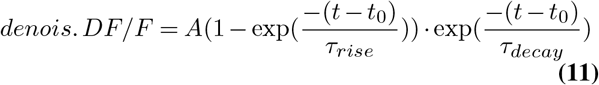

with *τ*_*rise*_ = 130 ms and *τ*_*decay*_ = 510 ms the rise and decay time of the calcium indicator GCamp6s and *t*_0_ the time point of spike occurrence and the amplitude scale parameter A, which we set to a constant value. To yield a signal-to-noise ratio (SNR) similar to experimental data, we added Gaussian noise to the traces with a SNR similar to what we observe in real data. We calculated the fluorescent traces with noise as:

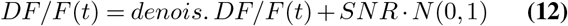

Finally, we subsampled the traces from the original temporal resolution of 1 kHz to the target frame rate 31 Hz by selecting the center data point of each time interval. This last step generated data with the same sampling rate as the two-photon microscopy data. From these resulting traces we then again obtained event rates using the deconvolution algorithm from the CaImAn package (Giovannucci et al., 2019). We applied the Bayesian decoding and found no difference in the decoding error for the slow and the fast session.

## Supplementary Figures

**Figure S1.**
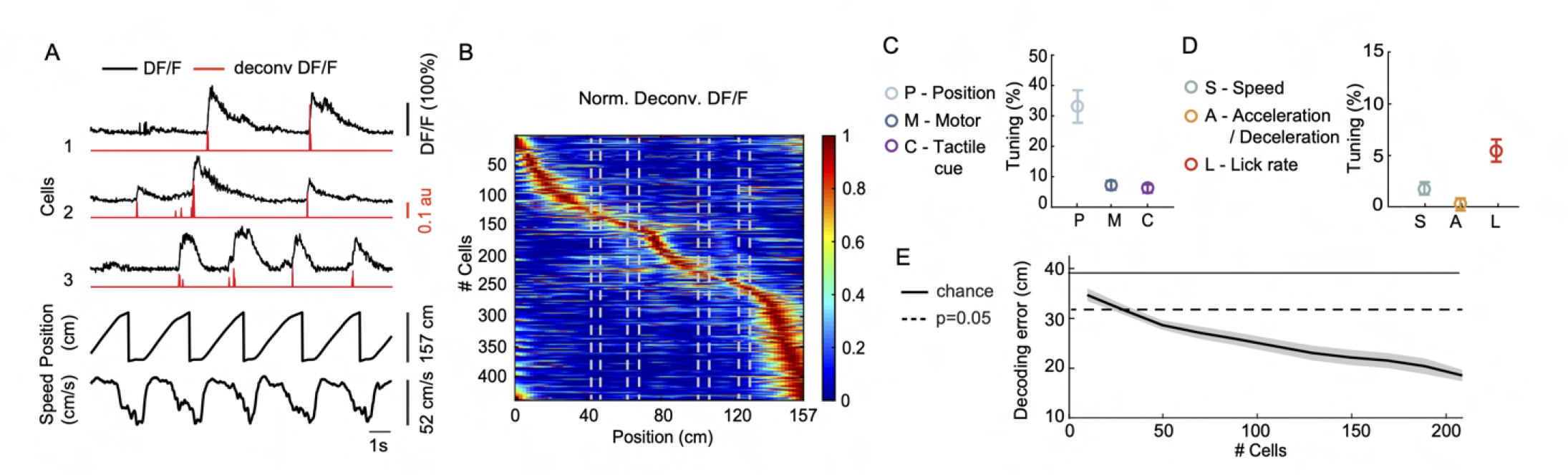
Position tuned neurons in L2/3 of agranular RSC. (**A**) Activity (DF/F) of three example RSC neurons across five laps, aligned to position and running speed. Inferred spike rate (deconvolved DF/F) shown in red. (**B**) Session-averaged activity of all position tuned RSC neurons sorted by peak activity. Dashed lines indicate tactile cue positions. (**C**) Percentage of RSC neurons tuned to position, motor variables and tactile cues (mean ± SEM across 9 sessions, 3 mice). Motor variables include running speed, acceleration/deceleration, and lick rate. (**D**) Percentage of RSC neurons tuned to speed, acceleration/deceleration, and lick rate (mean ± SEM across 9 sessions, 3 mice). (**D**) Decoding error (mean ± SEM across sessions) as a function of the number of RSC neurons.

**Figure S2.**
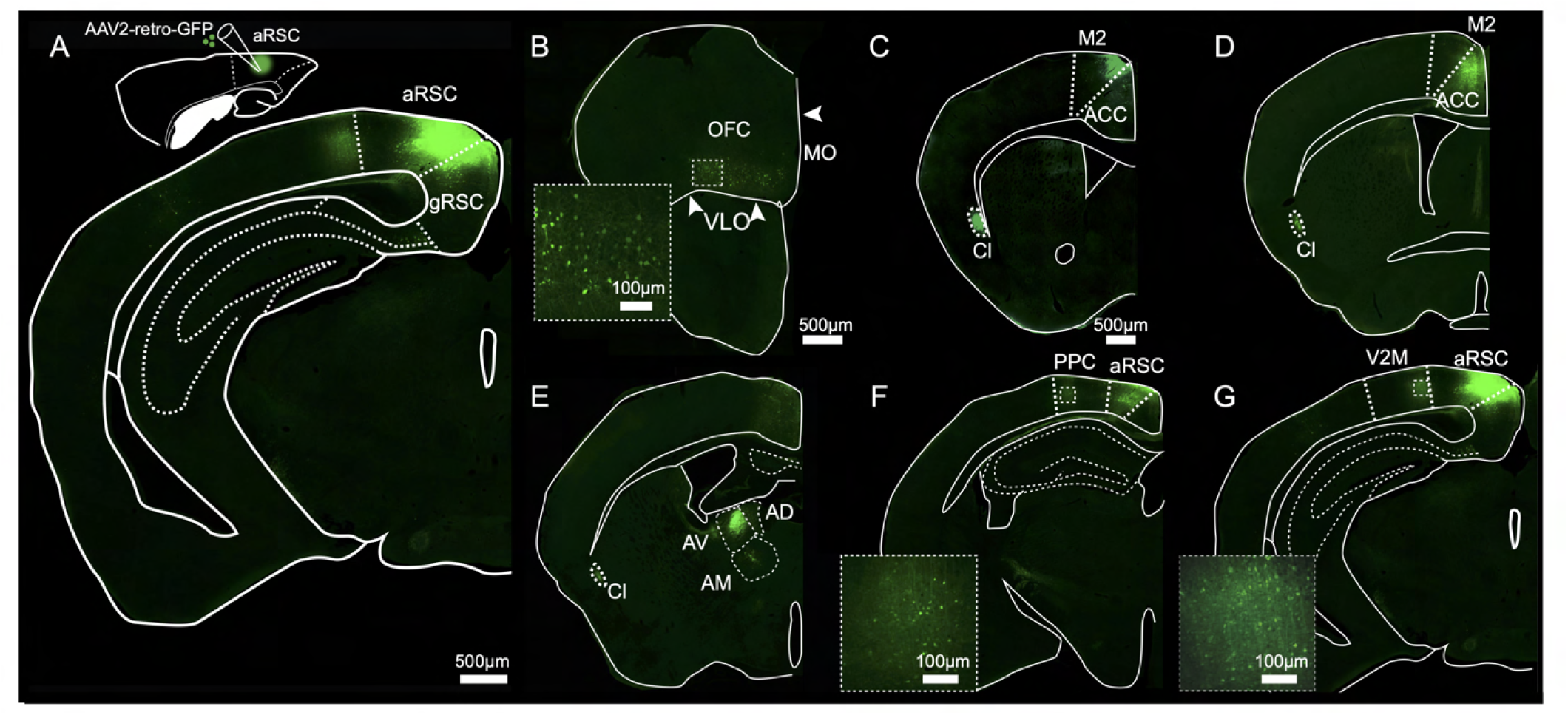
Retrograde viral labelling of neurons projecting to agranular RSC. (**A**) Top, sagittal mouse brain section showing injection of AAV-retro-GFP in agranular (aRSC). Bottom, coronal section showing the core of the injection site in aRSC. Delineation of brain areas was performed by superimposing the images of fluorescence and Nissl staining of the same section (see Methods). gRSC, granular RSC. (**B**) Retrogradely labelled neurons in orbitofrontal cortex (OFC). Inset shows magnified GFP-labelled cell bodies. VLO, ventrolateral OFC; MO, medial OFC. (**C**) Retrogradely labelled neurons in secondary motor cortex (M2). ACC, anterior cingulate cortex; Cl, claustrum. (**D**) Retrogradely labelled neurons in ACC. (**E**) Retrogradely labelled neurons in thalamus. AD, anterodorsal; AM, anteromedial; AV, anteroventral. (**F**) Retrogradely labelled neurons in posterior parietal cortex (PPC). Inset shows magnified cell bodies. (**G**) Retrogradely labelled neurons in medial secondary visual cortex (V2M). Inset shows magnified cell bodies. Scalebar from C also applies to D-G.

**Figure S3.**
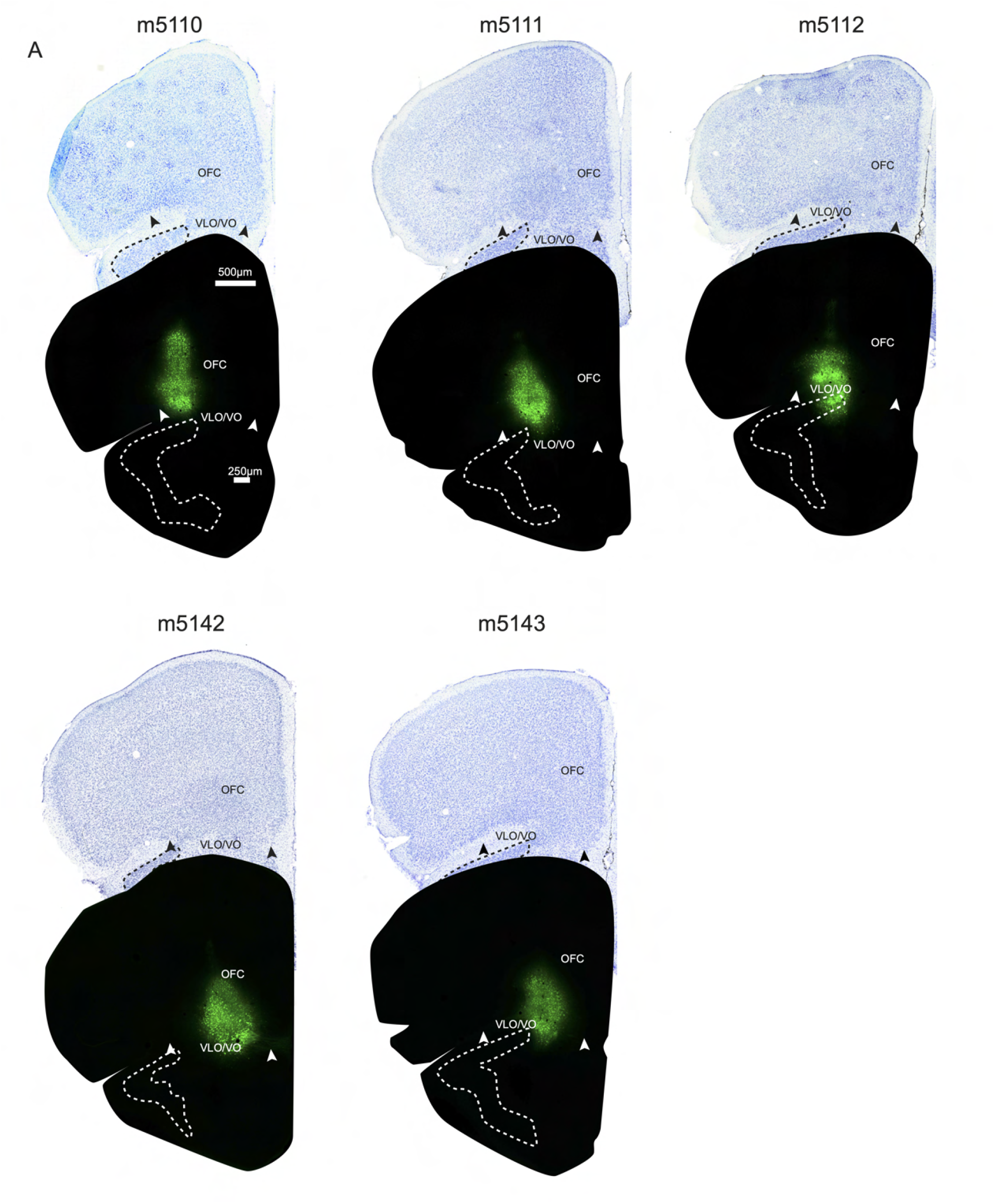
Delineation of injection sites in orbitofrontal cortex. (**A**) AAV-syn-GCaMP6s injection sites in OFC. Aligned fluorescence microscope image (bottom) showing the core of the injection, and bright field image of the same brain section stained with Nissl (top). The Nissl staining was used for atlas registration. Each panel represents one mouse.

**Figure S4.**
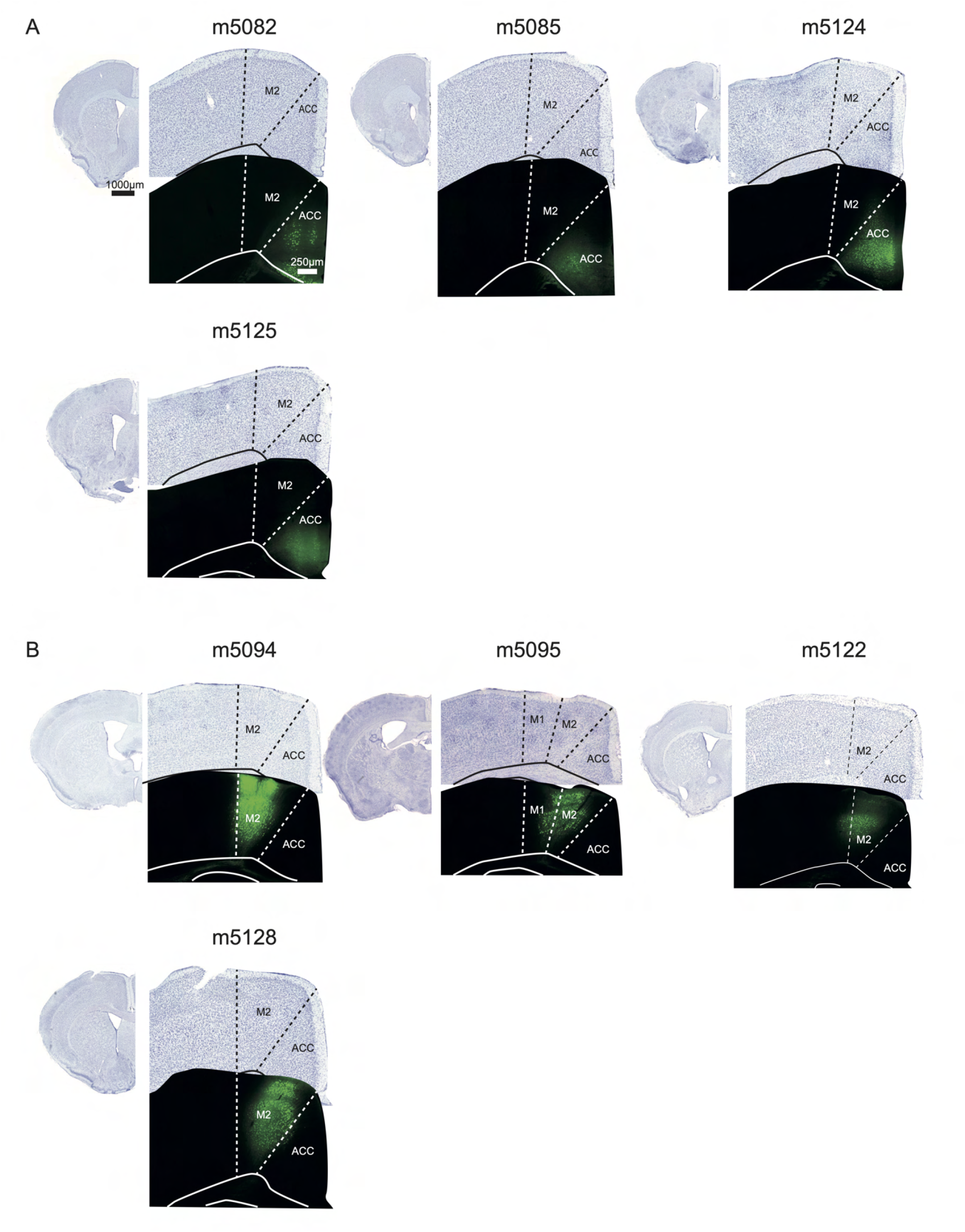
Delineating of injection sites in anterior cingulate cortex and secondary motor cortex. (**A**) AAV-syn-GCaMP6s injection sites in anterior cingulate cortex (ACC). On the left of each panel: hemispheric overview of the brain sections on the right. Aligned fluorescence microscope image (bottom) showing the core of the injection, and bright field image of the same brain section stained with Nissl (top). The Nissl staining was used for atlas registration. Each panel represents one mouse. (**B**) Same for injection sites in secondary motor cortex (M2).

**Figure S5.**
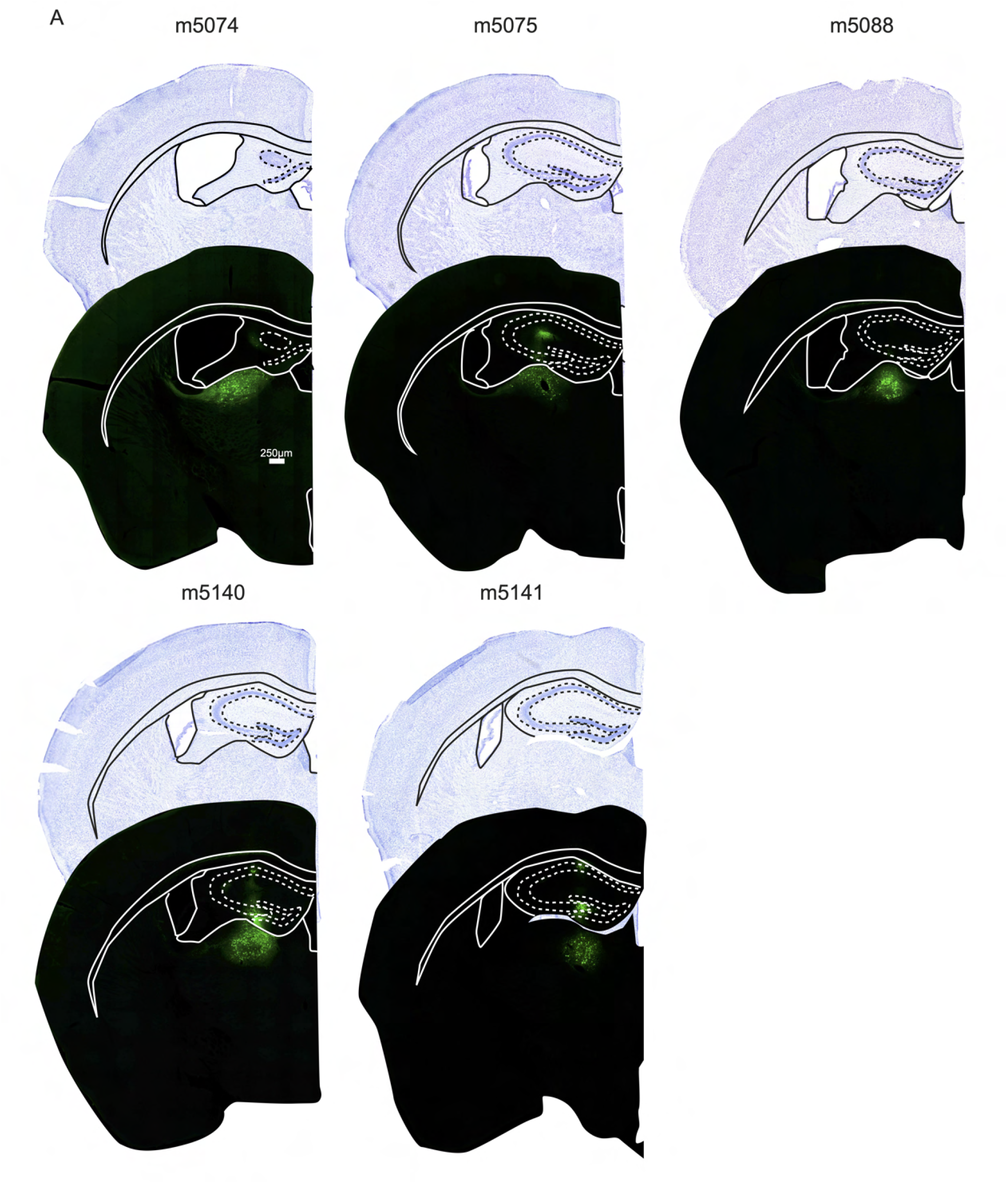
Delineation of injection sites in thalamus. (**A**) AAV-syn-GCaMP6s injection sites in thalamus. Aligned fluorescence microscope image (bottom) showing the core of the injection, and bright field image of the same brain section stained with Nissl (top). Each panel represents one mouse.

**Figure S6.**
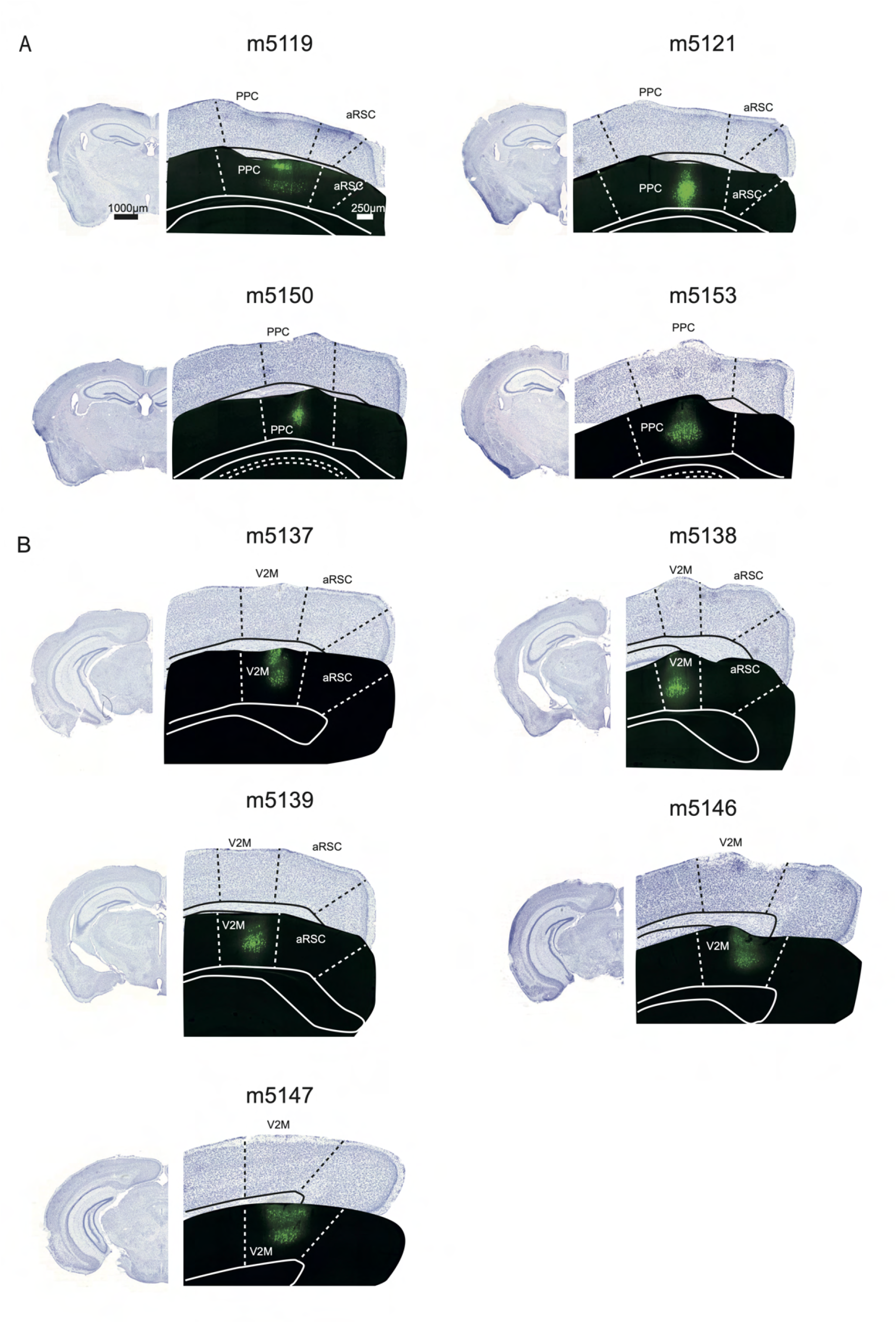
Delineation of injection sites in posterior parietal cortex and medial secondary visual cortex. (**A**) AAV-syn-GCaMP6s injection sites in posterior parietal cortex (PPC). On the left of each panel: hemispheric overview of the brain sections on the right. Aligned fluorescence microscope image (bottom) showing the core of the injection, and bright field image of the same brain section stained with Nissl (top). The Nissl staining was used for atlas registration. Each panel represents one mouse. (**B**) Same for injection sites in the medial secondary visual cortex (V2M).

**Figure S7.**
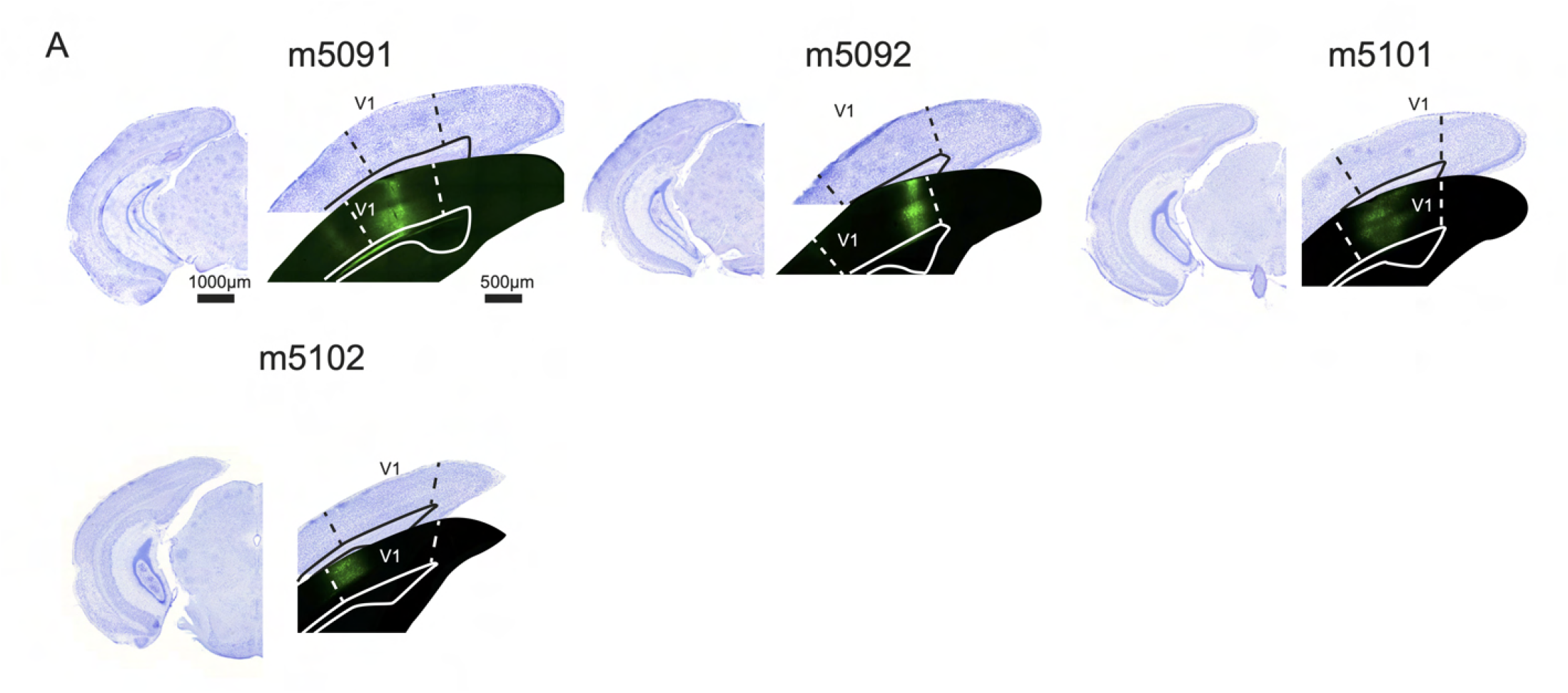
Delineation of injection sites in primary visual cortex. (**A**) AAV-syn-GCaMP6s injection sites in primary visual cortex (V1). On the left of each panel: hemispheric overview of the brain sections on the right. Aligned fluorescence microscope image (bottom) showing the core of the injection, and bright field image of the same brain section stained with Nissl (top). The Nissl staining was used for atlas registration. Each panel represents one mouse.

**Figure S8.**
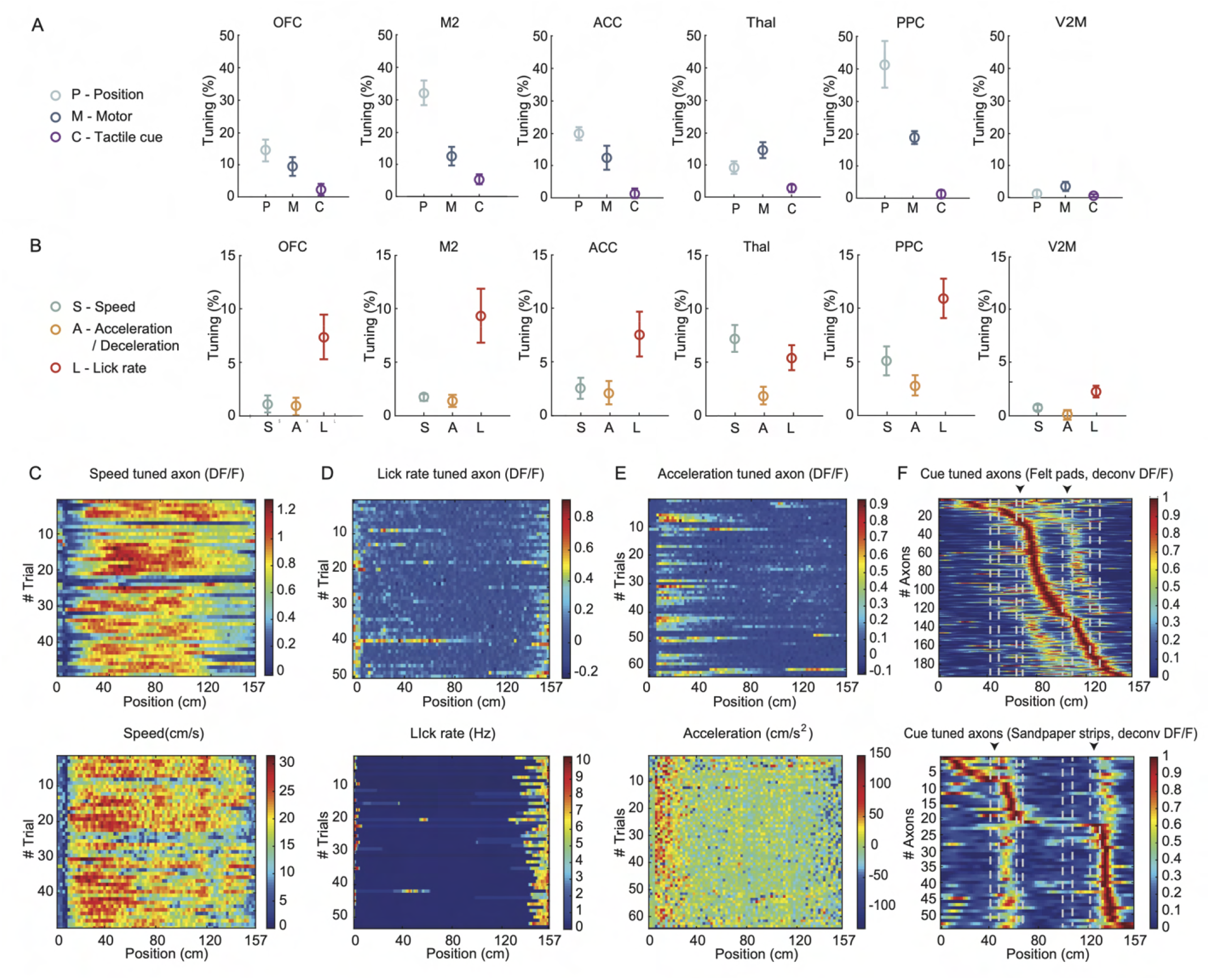
Summary of tuning properties of all long-range projections in RSC. (**A**) Percentage of neurons tuned to position, motor variables and tactile cues (mean ± SEM across sessions). Motor variables include running speed, acceleration and deceleration, and lick rate. (**B**) Percentage of neurons tuned to speed, acceleration/deceleration, and lick rate (mean ± SEM across sessions). (**C**) Top, activity (DF/F) of an example axon classified as speed tuned. Bottom, running speed of the mouse in the same session. (**D**) Top, activity (DF/F) of an example axon classified as tuned to lick rate. Bottom, lick rate of the mouse in the same session. (**E**) Top, activity (DF/F) of an example axon tuned to acceleration. Bottom, acceleration of the mouse in the same session. (**F**) Session-averaged activity (DF/F) of axons classified as tuned to the tactile cues; Top, axons tuned to the felt pads (only M2 axons shown). Bottom, axons tuned to the sandpaper strips (only thalamus axons shown). Axons are sorted by the location of their peak activity. Dashed lines indicate the positions of the cues: The inner pair of cues are felt pads; the outer pair are sandpaper strips.

**Figure S9.**
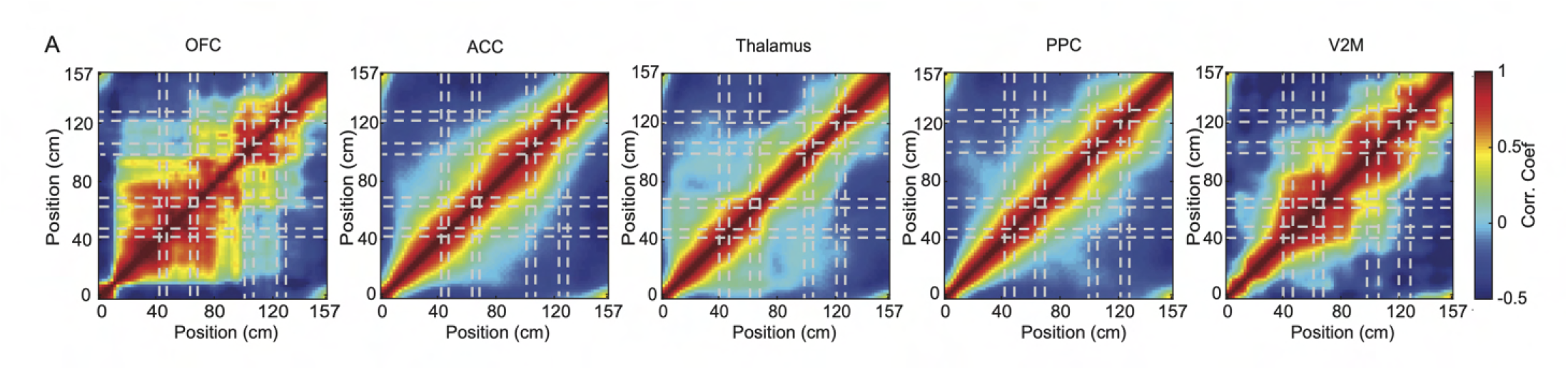
Correlations of population activity vectors of all long-range projections. (**A**) Correlation matrices (Pearson coefficient) of population activity vectors for all position tuned axons. The dashed lines indicate the position of the tactile cues.

**Figure S10.**
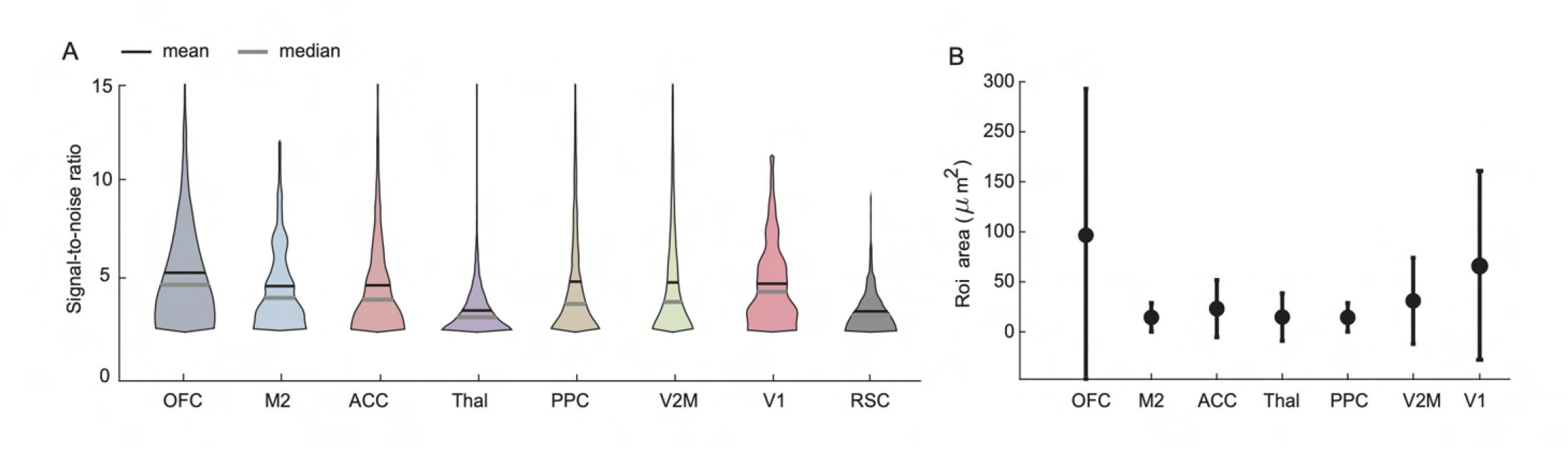
Signal to noise ratio, and region of interest (ROI) size for each projection. (**A**) Signal-to-Noise ratio distributions of axonal fluorescence signals (mean, black median, grey) for all axons. RSC soma data shown for comparison. (**B**) Size of the axonal regions of interest after image segmentation (mean ± SD across axons).

**Figure S11.**
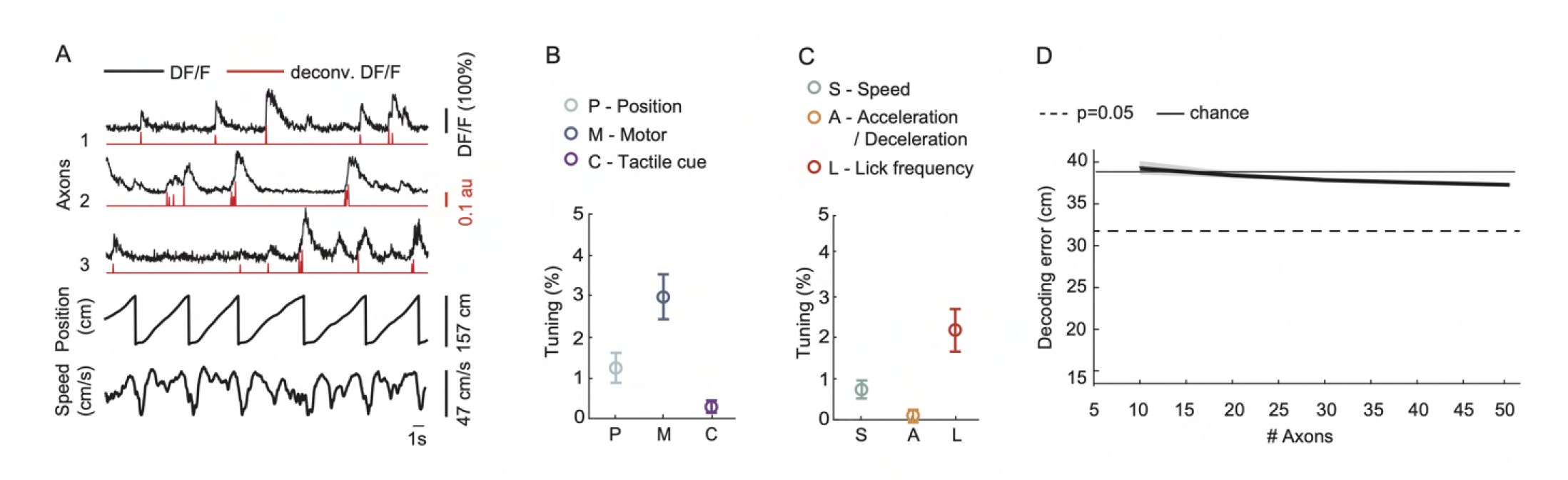
Functional imaging of axons from primary visual cortex projecting to posterior agranular RSC. (**A**) Activity (DF/F) of three example primary visual cortex (V1) axons across six laps, aligned to position and running speed. Inferred spike rate (deconvolved DF/F) shown in red. (**B**) Percentage of V1 axons tuned to position, motor variables and tactile cues (mean ± SEM across 10 sessions, 244 axons, 4 mice). Motor variables include running speed, acceleration/deceleration, and lick rate. (**C**) Percentage of V1 axons tuned to speed, acceleration/deceleration, and lick rate (mean ± SEM across 10 sessions, 4 mice). (**D**) Decoding error (mean ± SEM across sessions) as a function of number of V1 axons.

**Figure S12.**
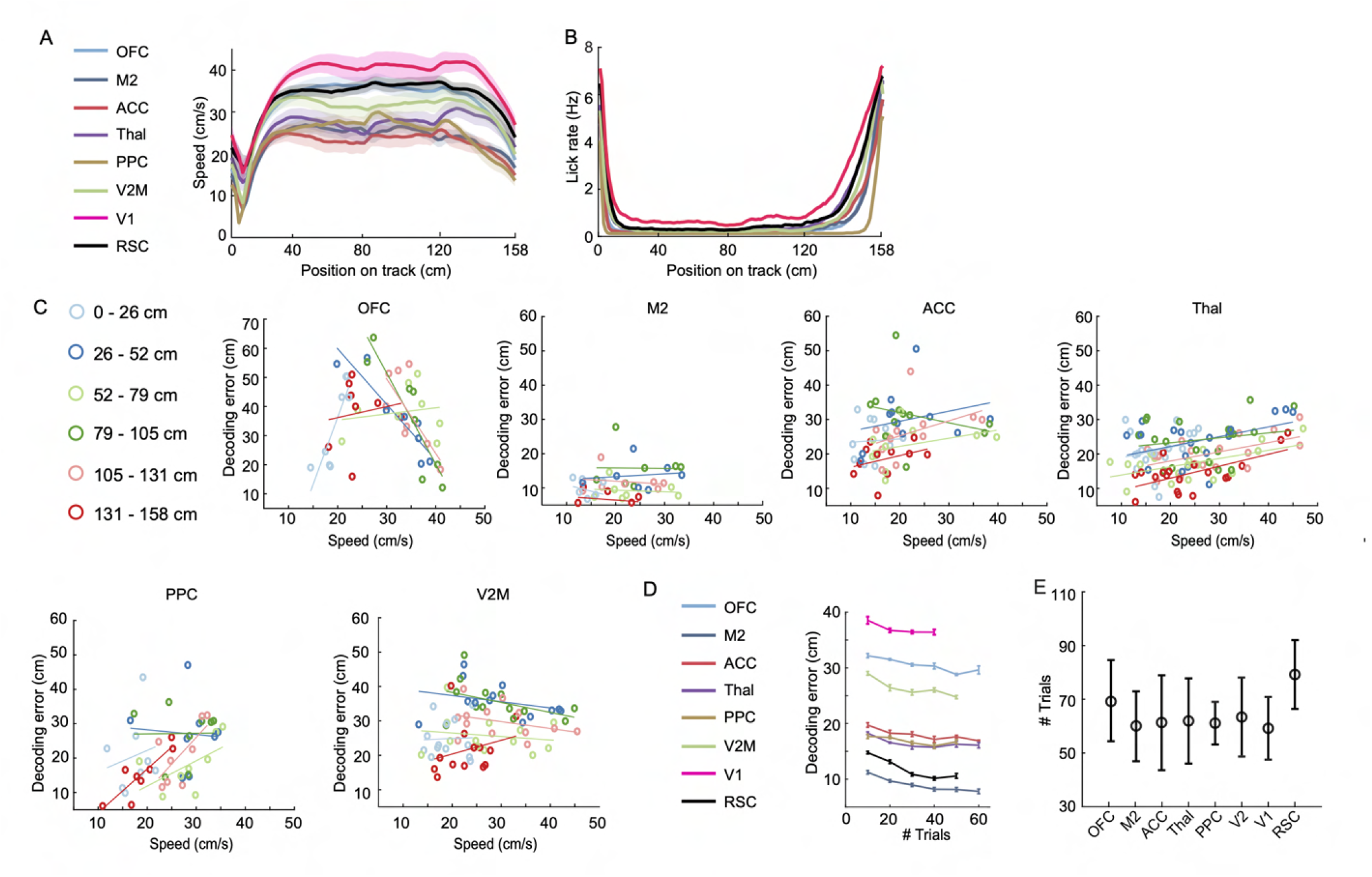
Analysis of relationship between running speed, number of trials per session and position decoding error. (**A**) Running speed all of mice (mean ± SEM across sessions). (**B**) Lick rate of all mice (mean across sessions). (**C**) Mean decoding error as a function of running speed. The track was divided in seven color-coded segments and for each segment the decoding error and average speed was calculated (each data point corresponds to one session). A linear fit is applied to all data points for a specific track segment. (**D**) Decoding error as a function of number of trials (mean ± SEM across cross-validation folds, one session per area). (**E**) Number of all trials performed by the mice (mean ± SD across sessions).

**Figure S13.**
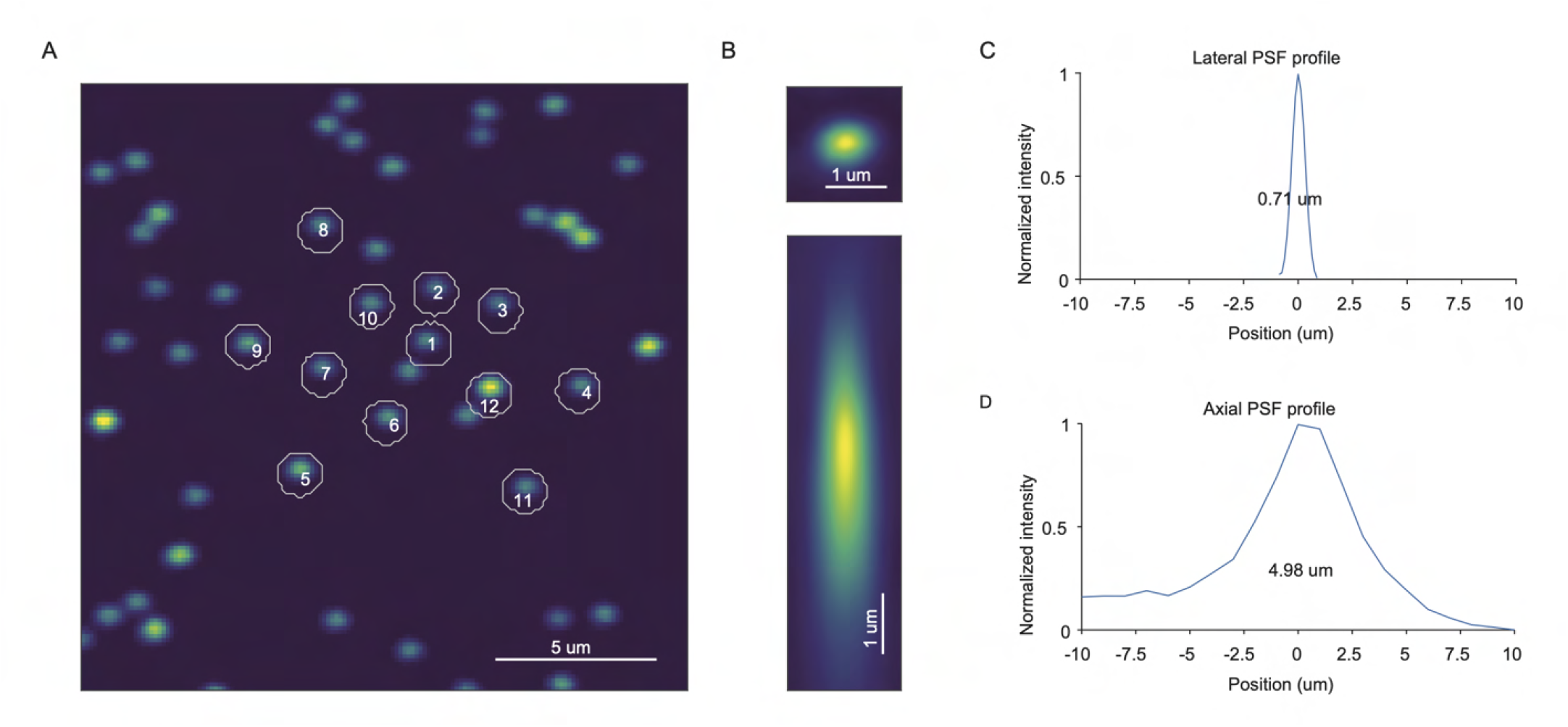
Underfilling the objective back aperture to enhance the axial point spread function during two-photon microscopy. (**A**) Two-photon fluorescence image of 175 nm fluorescent beads mounted on a microscope glass. The numbered beads were used to measure the point spread function (PSF). (**B**) Images of beads acquired using the optical system in the focal plane (top, x-y plane) and in a plane comprising the optical axis (bottom, y-z plane). (**C**,**D**) Average fluorescence profile of the images in (B). The full width at half maximum of the PSF is indicated.

**Figure S14.**
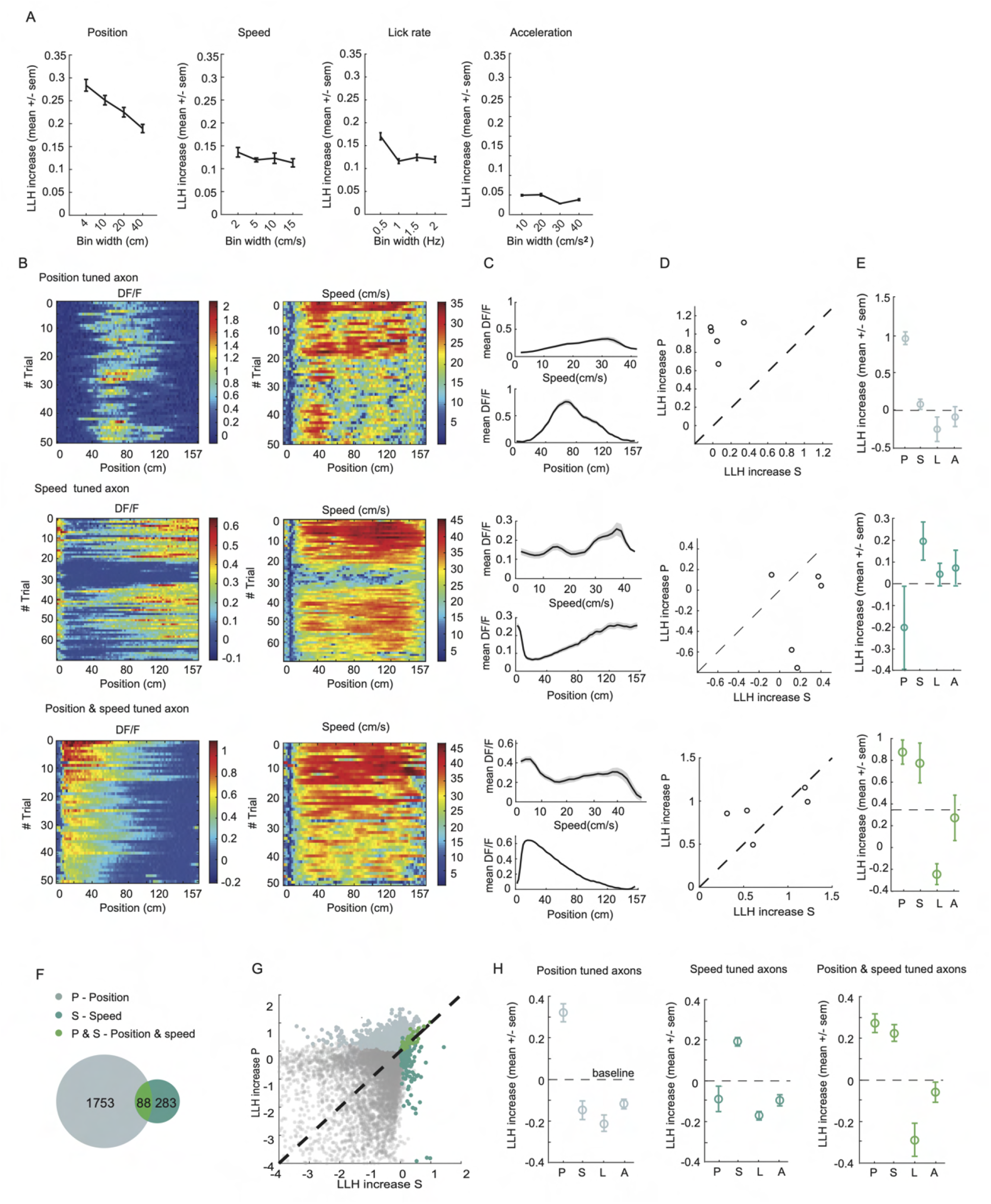
Examples of model performance for classifying position and speed. (**A**) Model performance as a function of bin width for position, speed, lick rate and acceleration for an example session (mean ± SEM log-likelihood increase over axons that performed better than chance for the variable). (**B**) (Left column) Example activity (DF/F) of three example axons classified as position-tuned (top), running speed-tuned (middle), and both position- and speed-tuned (bottom). (Right column) Running speed of the sessions corresponding to the data in the left column. (**C**) Tuning curves (averaged DF/F, same data as in (B)) over speed and position (mean ± SEM over trials) for the axons that were classified as position tuned (top), speed tuned (middle) or position and speed tuned (bottom). (**D**) Model performance comparison (LLH increase) of the speed model versus the position model (same data as in (B)). (**A**) Model performance across all models for the single axons shown in (E)) classified as position tuned (top), speed tuned (middle) or position and speed tuned (bottom) (mean ± SEM log-likelihood increase across cross validation folds). (**F**) Venn diagram that describes the quantities of axons that are characterized as position, speed or position and speed modulated (27 mice, 70 sessions, 1752 position tuned cells, 283 speed tuned cells, 88 position and speed tuned cells, 10852 untuned cells, data from all long-range projections). (**G**) Position and speed model performances of axons that are classified as position, speed or position and speed modulated, and, in grey, of axons that are classified as not modulated to any of the variables. Each data point is the mean log-likelihood increase over cross validation folds (same data as in (F)). (**H**) Model performance across all models for all axons that are characterized as position, speed or position and speed modulated (mean ± SEM across areas, 27 mice, 70 sessions).

## Bibliography

Ahmed, O. J. and Mehta, M. R. (2009). The hippocampal rate code: anatomy, physiology and theory. Trends in Neurosciences, 32(6):329–338.

Alexander, A. S. and Nitz, D. A. (2015). Retrosplenial cortex maps the conjunction of internal and external spaces. Nature Neuroscience, 18(8):1143--1151.

Broussard, G. J. and Petreanu, L. (2021). Eavesdropping wires: recording activity in axons using genetically encoded calcium indicators. Journal of Neuroscience Methods, 360:109251.

Campbell, M. G., Attinger, A., Ocko, S. A., Ganguli, S., and Giocomo, L. M. (2021). Distance-tuned neurons drive specialized path integration calculations in medial entorhinal cortex. Cell Reports, 36(10):109669.

Cembrowski, M. S., Phillips, M. G., DiLisio, S. F., Shields, B. C., Winnubst, J., Chandrashekar, J., Bas, E., and Spruston, N. (2018). Dissociable Structural and Functional Hippocampal Outputs via Distinct Subiculum Cell Classes. Cell, pages 1–32.

Cenquizca, L. A. and Swanson, L. W. (2007). Spatial organization of direct hippocampal field CA1 axonal projections to the rest of the cerebral cortex. Brain Research Reviews, 56(1):1–26.

Cho, J. and Sharp, P. E. (2001). Head direction, place, and movement correlates for cells in the rat retrosplenial cortex. Behavioral Neuroscience, 115(1):3–25.

Cowansage, K. K., Shuman, T., Dillingham, B. C., Chang, A., Golshani, P., and Mayford, M. (2014). Direct reactivation of a coherent neocortical memory of context. Neuron, 84(2):432–441.

Czajkowski, R., Jayaprakash, B., Wiltgen, B., Rogerson, T., Guzman-Karlsson, M. C., Barth, A. L., Trachtenberg, J. T., and Silva, A. J. (2014). Encoding and storage of spatial information in the retrosplenial cortex. Proceedings of the National Academy of Sciences, 111(23):8661–8666.

Dana, H., Chen, T.-W., Hu, A., Shields, B. C., Guo, C., Looger, L. L., Kim, D. S., and Svoboda, K. (2014). Thy1-GCaMP6 Transgenic Mice for Neuronal Population Imaging In Vivo. PLoS ONE, 9(9):e108697.

Diamanti, E. M., Reddy, C. B., Schröder, S., Muzzu, T., Harris, K. D., Saleem, A. B., and Carandini, M. (2021). Spatial modulation of visual responses arises in cortex with active navigation. eLife, 10:e63705.

Eichenbaum, H. (2017). The role of the hippocampus in navigation is memory. Journal of neurophysiology, 117(4):1785–1796.

Esteves, I. M., Chang, H., Neumann, A. R., Sun, J., Mohajerani, M. H., and Mc-Naughton, B. L. (2020). Spatial Information Encoding Across Multiple Neocortical Regions Depends on an Intact Hippocampus. Journal of Neuroscience, 41(2):JN–RM–1788–20.

Fischer, L. F., Soto-Albors, R. M., Buck, F., and Harnett, M. T. (2020). Representation of visual landmarks in retrosplenial cortex. eLife, 9:e51458.

Fiser, A., Mahringer, D., Oyibo, H. K., Petersen, A. V., Leinweber, M., and Keller, G. B. (2016). Experience-dependent spatial expectations in mouse visual cortex. Nature Neuroscience, 19(12):1658–1664.

Flossmann, T. and Rochefort, N. L. (2021). Spatial navigation signals in rodent visual cortex. Current Opinion in Neurobiology, 67:163–173.

Fournier, J., Saleem, A. B., Diamanti, E. M., Wells, M. J., Harris, K. D., and Carandini, M. (2020). Mouse Visual Cortex Is Modulated by Distance Traveled and by Theta Oscillations. Current Biology, 30(19):3811–3817.e6.

Franco, L. M. and Goard, M. J. (2021). A distributed circuit for associating environmental context with motor choice in retrosplenial cortex. Science Advances, 7(35):eabf9815.

Friedrich, J., Zhou, P., and Paninski, L. (2017). Fast online deconvolution of calcium imaging data. PLOS Computational Biology, 13(3):e1005423.

Giovannucci, A., Friedrich, J., Gunn, P., Kalfon, J., Brown, B. L., Koay, S. A., Taxidis, J., Najafi, F., Gauthier, J. L., Zhou, P., Khakh, B. S., Tank, D. W., Chklovskii, D. B., and Pnevmatikakis, E. A. (2019). CaImAn an open source tool for scalable calcium imaging data analysis. eLife, 8:e38173.

Grieves, R. M. and Jeffery, K. J. (2016). The representation of space in the brain. Behavioural Processes, 135:113–131.

Groen, T. v. and Wyss, J. M. (1992). Connections of the retrosplenial dysgranular cortex in the rat. The Journal of comparative neurology, 315(2):200–216.

Guo, Z. V., Hires, S. A., Li, N., O’Connor, D. H., Komiyama, T., Ophir, E., Huber, D., Bonardi, C., Morandell, K., Gutnisky, D., Peron, S., Xu, N.-l., Cox, J., and Svoboda, K. (2014). Procedures for Behavioral Experiments in Head-Fixed Mice. PLoS ONE, 9(2):e88678.

Haggerty, D. C. and Ji, D. (2015). Activities of visual cortical and hippocampal neurons co-fluctuate in freely moving rats during spatial behavior. eLife, 4:e08902.

Hardcastle, K., Maheswaranathan, N., Ganguli, S., and Giocomo, L. M. (2017). A Multiplexed, Heterogeneous, and Adaptive Code for Navigation in Medial Entorhinal Cortex. Neuron, 94(2):375–387.e7.

Harvey, C. D., Coen, P., and Tank, D. W. (2012). Choice-specific sequences in parietal cortex during a virtual-navigation decision task. Nature, 484(7392):62–68.

Haugland, K. G., Sugar, J., and Witter, M. P. (2019). Development and topographical organization of projections from the hippocampus and parahippocampus to the retrosplenial cortex. The European Journal of Neuroscience, 50(1):1799–1819.

Hok, V., Save, E., Lenck-Santini, P. P., and Poucet, B. (2005). Coding for spatial goals in the prelimbic/infralimbic area of the rat frontal cortex. Proceedings of the National Academy of Sciences of the United States of America, 102(12):4602–4607.

Holtmaat, A., Bonhoeffer, T., Chow, D. K., Chuckowree, J., Paola, V. D., Hofer, S. B., Hübener, M., Keck, T., Knott, G., Lee, W.-C. A., Mostany, R., Mrsic-Flogel, T. D., Nedivi, E., Portera-Cailliau, C., Svoboda, K., Trachtenberg, J. T., and Wilbrecht, L. (2009). Long-term, high-resolution imaging in the mouse neocortex through a chronic cranial window. Nature Protocols, 4(8):1128–1144.

Hovde, K., Gianatti, M., Witter, M. P., and Whitlock, J. R. (2018). Architecture and organization of mouse posterior parietal cortex relative to extrastriate areas. European Journal of Neuroscience, 49(10):1313–1329.

Jankowski, M. M. and O’Mara, S. M. (2015). Dynamics of place, boundary and object encoding in rat anterior claustrum. Frontiers in Behavioral Neuroscience, 9:250.

Jankowski, M. M., Passecker, J., Islam, M. N., Vann, S., Erichsen, J. T., Aggleton, J. P., and O’Mara, S. M. (2015). Evidence for spatially-responsive neurons in the rostral thalamus. Frontiers in behavioral neuroscience, 9:425–18.

Jay, T. M. and Witter, M. P. (1991). Distribution of hippocampal CA1 and subicular efferents in the prefrontal cortex of the rat studied by means of anterograde transport of Phaseolus vulgaris-leucoagglutinin. Journal of Comparative Neurology, 313(4):574–586.

Kitanishi, T., Umaba, R., and Mizuseki, K. (2021). Robust information routing by dorsal subiculum neurons. Science Advances, 7(11):eabf1913.

Kjelstrup, K. B., Solstad, T., Brun, V. H., Hafting, T., Leutgeb, S., Witter, M. P., Moser, E. I., and Moser, M.-B. (2008). Finite Scale of Spatial Representation in the Hippocampus. Science, 321(5885):140–143.

Knierim, J. J. (2006). Neural representations of location outside the hippocampus. Learning & Memory, 13(4):405–415.

Long, X., Deng, B., Cai, J., Chen, Z. S., and Zhang, S.-J. (2021). A compact spatial map in V2 visual cortex. bioRxiv, page 2021.02.11.430687.

Long, X. and Zhang, S.-J. (2021). A novel somatosensory spatial navigation system outside the hippocampal formation. Cell Research, 31(6):649–663.

Lütcke, H., Gerhard, F., Zenke, F., Gerstner, W., and Helmchen, F. (2013). Inference of neuronal network spike dynamics and topology from calcium imaging data. Frontiers in neural circuits, 7(December):201.

Mao, D., Kandler, S., McNaughton, B. L., and Bonin, V. (2017). Sparse orthogonal population representation of spatial context in the retrosplenial cortex. Nature communications, pages 1–9.

Mao, D., Molina, L. A., Bonin, V., and McNaughton, B. L. (2020). Vision and Locomotion Combine to Drive Path Integration Sequences in Mouse Retrosplenial Cortex. Current Biology.

Mao, D., Neumann, A. R., Sun, J., Bonin, V., Mohajerani, M. H., and McNaughton, B. L. (2018). Hippocampus-dependent emergence of spatial sequence coding in retrosplenial cortex. Proceedings of the National Academy of Sciences, 115(31):8015–8018.

McNaughton, B. L., Battaglia, F. P., Jensen, O., Moser, E. I., and Moser, M.-B. (2006). Path integration and the neural basis of the ‘cognitive map’. Nature reviews. Neuroscience, 7(8):663–678.

McNaughton, B. L., Mizumori, S. J. Y., Barnes, C. a., Leonard, B. J., Marquis, M., and Green, E. J. (1994). Cortical Representation of Motion during Unrestrained Spatial Navigation in the Rat. Cerebral Cortex, 4(1):27–39.

Milczarek, M. M., Vann, S. D., and Sengpiel, F. (2018). Spatial Memory Engram in the Mouse Retrosplenial Cortex. Current Biology, 28(12):1–13.

Mizumori, S., Ward, K., and Lavoie, A. (1992). Medial septal modulation of entorhinal single unit activity in anesthetized and freely moving rats. Brain Research, 570(1-2):188–197.

Murtagh, F. and Contreras, P. (2012). Algorithms for hierarchical clustering: an overview. Wiley Interdisciplinary Reviews: Data Mining and Knowledge Discovery, 2(1):86–97.

Nitz, D. a. (2006). Tracking Route Progression in the Posterior Parietal Cortex. Neuron, 49(5):747–756.

Nitzan, N., McKenzie, S., Beed, P., English, D. F., Oldani, S., Tukker, J. J., Buzsáki, G., and Schmitz, D. (2020). Propagation of hippocampal ripples to the neocortex by way of a subiculum-retrosplenial pathway. Nature Communications, 11(1):1947.

O’Keefe, J. and Dostrovsky, J. (1971). The hippocampus as a spatial map. Preliminary evidence from unit activity in the freely-moving rat. Brain Research, 34(1):171–175.

Olson, J. M., Li, J. K., Montgomery, S. E., and Nitz, D. A. (2020). Secondary Motor Cortex Transforms Spatial Information into Planned Action during Navigation. Current Biology, 30(10):1845–1854.e4.

Pakan, J. M. P., Currie, S. P., Fischer, L., and Rochefort, N. L. (2018). The Impact of Visual Cues, Reward, and Motor Feedback on the Representation of Behaviorally Relevant Spatial Locations in Primary Visual Cortex. Cell Reports, 24(10):2521–2528.

Paxinos, G. and Franklin, K. (2019). The Mouse in stereotaxic coordinates. Elsevier, 5th edition edition.

Petreanu, L., Gutnisky, D. A., Huber, D., Xu, N.-l., O’Connor, D. H., Tian, L., Looger, L., and Svoboda, K. (2012). Activity in motor-sensory projections reveals distributed coding in somatosensation. Nature, 489(7415):299–303.

Peyrache, A. and Duszkiewicz, A. J. (2021). A spatial map out of place. Cell Research, 31(6):605–606.

Pnevmatikakis, E. a., Soudry, D., Gao, Y., Machado, T. A., Merel, J., Pfau, D., Reardon, T., Mu, Y., Lacefield, C., Yang, W., Ahrens, M., Bruno, R., Jessell, T. M., Peterka, D. S., Yuste, R., and Paninski, L. (2016). Simultaneous Denoising, Deconvolution, and Demixing of Calcium Imaging Data. Neuron, 89(2):285–299.

Powell, A., Connelly, W. M., Vasalauskaite, A., Nelson, A. J. D., Vann, S. D., Aggleton, J. P., Sengpiel, F., and Ranson, A. (2020). Stable Encoding of Visual Cues in the Mouse Retrosplenial Cortex. Cerebral Cortex (New York, NY), 30(8):4424–4437.

Quirk, G., Muller, R., Kubie, J., and Ranck, J. (1992). The positional firing properties of medial entorhinal neurons: description and comparison with hippocampal place cells. The Journal of Neuroscience, 12(5):1945–1963.

Rajan, K., Harvey, C. D., and Tank, D. W. (2016). Recurrent Network Models of Sequence Generation and Memory. Neuron, 90(1):128–142.

Remondes, M. and Wilson, M. A. (2013). Cingulate-Hippocampus Coherence and Trajectory Coding in a Sequential Choice Task. Neuron, 80(5):1277–1289.

Robinson, N. T., Descamps, L. A., Russell, L. E., Buchholz, M. O., Bicknell, B. A., Antonov, G. K., Lau, J. Y., Nutbrown, R., Schmidt-Hieber, C., and Häusser, M. (2020). Targeted Activation of Hippocampal Place Cells Drives Memory-Guided Spatial Behavior. Cell, 183(6):1586–1599.e10.

Royer, S., Zemelman, B. V., Losonczy, A., Kim, J., Chance, F., Magee, J. C., and Buzsáki, G. (2012). Control of timing, rate and bursts of hippocampal place cells by dendritic and somatic inhibition. Nature neuroscience, 15(5):769–775.

Saleem, A. B., Diamanti, E. M., Fournier, J., Harris, K. D., and Carandini, M. (2018). Coherent encoding of subjective spatial position in visual cortex and hippocampus. Nature, 562(7725).

Skelin, I., Kilianski, S., and McNaughton, B. L. (2019). Hippocampal coupling with cortical and subcortical structures in the context of memory consolidation. Neurobiology of Learning and Memory, 160:21–31.

Strange, B. A., Witter, M. P., Lein, E. S., and Moser, E. I. (2014). Functional organization of the hippocampal longitudinal axis. Nature Publishing Group, 15(10):655–669.

Sugar, J., Witter, M. P., Strien, N. M. v., and Cappaert, N. L. M. (2011). The retrosplenial cortex: intrinsic connectivity and connections with the (para)hippocampal region in the rat. An interactive connectome. Frontiers in Neuroinformatics, 5:7.

Tervo, D. G. R., Hwang, B.-Y., Viswanathan, S., Gaj, T., Lavzin, M., Ritola, K. D., Lindo, S., Michael, S., Kuleshova, E., Ojala, D., Huang, C.-C., Gerfen, C. R., Schiller, J., Dudman, J. T., Hantman, A. W., Looger, L. L., Schaffer, D. V., and Karpova, A. Y. (2016). A Designer AAV Variant Permits Efficient Retrograde Access to Projection Neurons. Neuron, 92(2):372–382.

Teyler, T. J. and DiScenna, P. (1986). The hippocampal memory indexing theory. Behavioral neuroscience, 100(2):147–154.

Vann, S. D. and Aggleton, J. P. (2002). Extensive cytotoxic lesions of the rat retrosplenial cortex revel consistent deficits on tasks that tax allocentric spatial memory. Behavioral neuroscience, 116(1):85–94.

Vann, S. D., Aggleton, J. P., and Maguire, E. A. (2009). What does the retrosplenial cortex do? Nature reviews. Neuroscience, 10(11):792–802.

Wang, C., Chen, X., and Knierim, J. J. (2020). Egocentric and allocentric representations of space in the rodent brain. Current Opinion in Neurobiology, 60:12–20.

Wyss, J. M. and Groen, T. v. (1992). Connections between the retrosplenial cortex and the hippocampal formation in the rat: a review. Hippocampus, 2(1):1–11.

Yamawaki, N., Radulovic, J., and Shepherd, G. M. G. (2016). A Corticocortical Circuit Directly Links Retrosplenial Cortex to M2 in the Mouse. The Journal of neuroscience : the official journal of the Society for Neuroscience, 36(36):9365–9374.

Zipfel, W. R., Williams, R. M., and Webb, W. W. (2003). Nonlinear magic: multiphoton microscopy in the biosciences. Nature Biotechnology, 21(11):1369–1377.

